# A conformational switch controlling the toxicity of the prion protein

**DOI:** 10.1101/2021.09.20.460912

**Authors:** Karl Frontzek, Marco Bardelli, Assunta Senatore, Anna Henzi, Regina R. Reimann, Seden Bedir, Marika Marino, Rohanah Hussain, Simon Jurt, Georg Meisl, Mattia Pedotti, Federica Mazzola, Giuliano Siligardi, Oliver Zerbe, Marco Losa, Tuomas Knowles, Asvin Lakkaraju, Caihong Zhu, Petra Schwarz, Simone Hornemann, Matthew G. Holt, Luca Simonelli, Luca Varani, Adriano Aguzzi

## Abstract

Prion infections cause conformational changes of PrP^C^ and lead to progressive neurological impairment. Here we show that toxic, prion-mimetic ligands induce an intramolecular R208-H140 hydrogen bond (“H-latch”) altering the flexibility of the α2-α3 and β2-α2 loops of PrP^C^. Expression of a PrP^2Cys^ mutant mimicking the H-latch was constitutively toxic, whereas a PrP^R207A^ mutant unable to form the H-latch conferred resistance to prion infection. High-affinity ligands that prevented H-latch induction repressed prion-related neurodegeneration in organotypic cerebellar cultures. We then selected phage-displayed ligands binding wild-type PrP^C^, but not PrP^2Cys^. These binders depopulated H-latched conformers and conferred protection against prion toxicity. Finally, brain-specific expression of an antibody rationally designed to prevent H-latch formation, prolonged the life of prion-infected mice despite unhampered prion propagation, confirming that the H-latch is causally linked to prion neurotoxicity.

## Main text

The neurotoxicity of prions requires the interaction of the misfolded prion protein PrP^Sc^ with its cellular counterpart PrP^C^, which ultimately leading to depletion of the PIKfyve kinase (*1*) and to spongiform encephalopathy. Prion toxicity is initiated by unknown mechanisms that require membrane-bound PrP^C^ (*2, 3*). Antibodies binding the globular domain (GD) of PrP^C^ can halt this process (*4*), but they can also activate toxic intracellular cascades (*5–7*). Similar events occur in prion-infected brains, and substances that counteract the damage of infectious prions can also alleviate the toxicity of anti-PrP^C^ antibodies such as POM1 (*6*). This suggests that POM1 and prions exert their toxicity through similar mechanisms. Structural analysis and molecular dynamics (MD) simulations indicated that POM1 induces an intramolecular hydrogen bond in both human and murine PrP^C^ between R208 and H139 in murine PrP^C^) (*8*). This “H-latch” constrains the POM1 epitope while allosterically increasing the flexibility of the β2-α2 and α2-α3 loops (Fig. 1, S1).

**Fig. 1.**
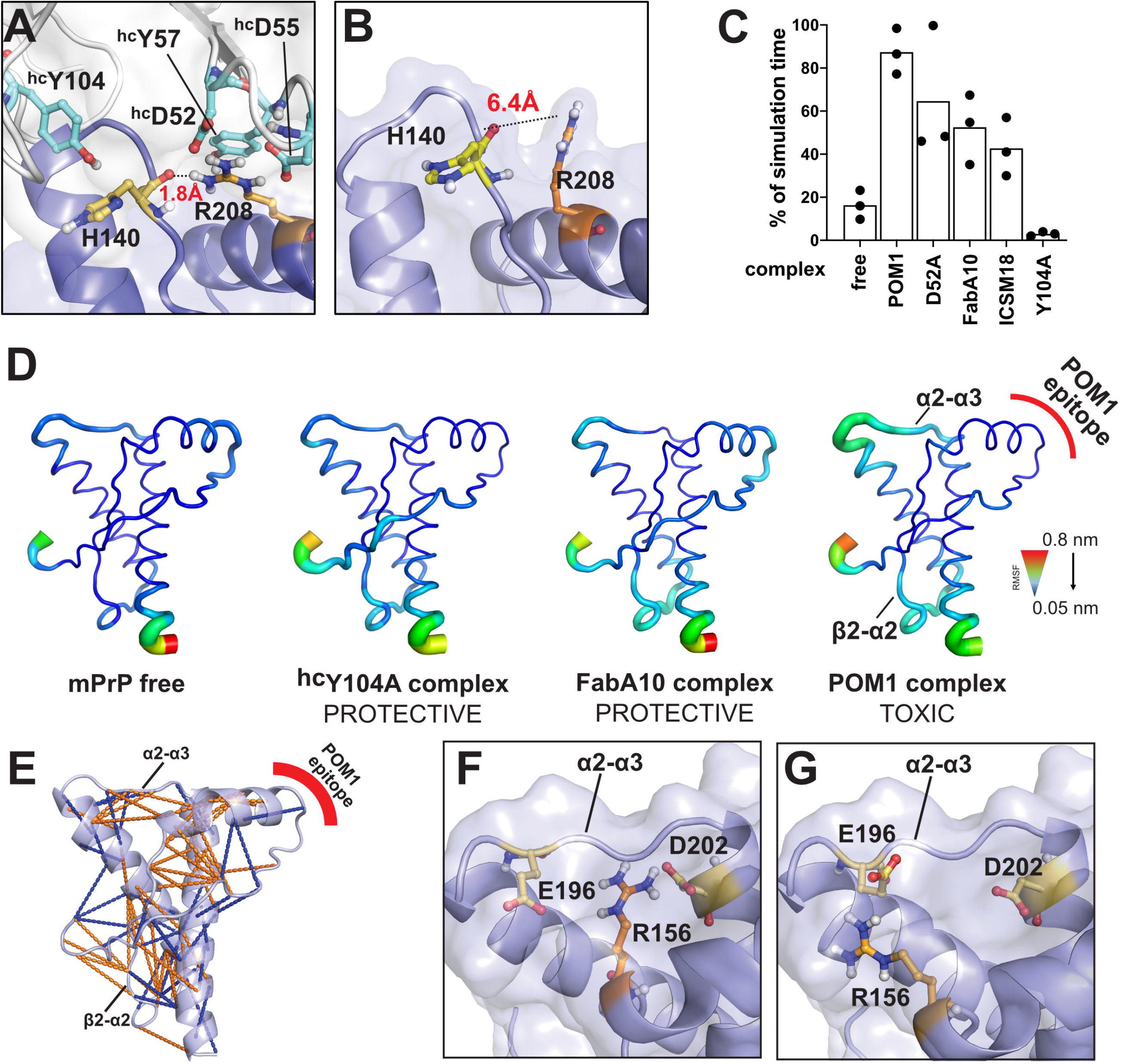
POM1 induces an intramolecular hydrogen bond between R208A and H140 of human PrP^c^. **(A)** Binding of PrP^C^ to the neurotoxic antibody POM1 favors formation of a R208-H140 hydrogen bond in the GD of PrP^C^ that is absent from free PrP^C^ **(B). (C)** Molecular dynamics simulations indicate that toxic antibodies are more likely to induce the R208-H140 bond. Ordinate: percentage of simulation time in which the H-bond is present. See also Fig. S1. **(D)** GD flexibility according to MD simulations. Narrow blue ribbons: rigidity; large green/red ribbons: increased flexibility. PrP bound to protective pomologs resembles free PrP. PrP bound to POM1 induces increased flexibility in the α2-α3 and β2-α2 loop. (E) Binding of the toxic Ab POM1 to PrP induces local structural changes within the GD, here shown as cartoon, both within and outside the epitope region. Side-chain contacts (less than 5Å) that are present only in PrP free (blue, PDB 1xyx) or PrP bound (orange, PDB 4H88) are indicated by lines. (F) POM1 binding breaks the R156-E196 interaction, increasing the α2-α3 flexibility, and induces the formation of a R156-D202 salt bridge. (G) R156 interacts with E196 in free PrP, which helps to rigidify the α2-α3 loop.

In order to explore its role in prion toxicity, we generated a murine PrP^R207A^ mutant that prevents the H-latch without altering the conformation of PrP (Fig. S1). We stably expressed mPrP^R207A^ in *Prnp*^-/-^ CAD5 cells (*9*) and *Prnp*^ZH3/ZH3^ cerebellar organotypic cultured slices (COCS, Fig. S2) (*10, 11*). A panel of conformation-specific anti-PrP antibodies showed similar staining patterns of PrP^C^ and mPrP^R207A^, confirming proper folding, except for reduced POM1 binding (Fig. S4A+B) as expected from the structure of PrP-POM1 co-crystals (*8*). *Prnp*^-/-^ CAD5 cells expressing mPrP^R207A^ were resistant to POM1 toxicity and, importantly, showed impaired prion replication (Fig. S3C-F), pointing to common toxic properties.

Lack of H-latch confers resistance to prion and POM1 toxicity. To test if its presence can induce toxicity even in the absence of ligands, we designed a R207C/I138C di-cysteine PrP^C^ mutant (PrP^2Cys^ Fig. S4) with the goal of replicating the structural effects of the H-latch in the absence of POM1 binding. NMR and MD analysis of recombinant mPrP^2Cys^ were consistent with a folded protein resembling the H-latch conformation (Fig. S4). PrP^2Cys^ expressed in a *Prnp*^-/-^ CAD5 cell line showed correct glycosylation and topology, and did not trigger unfolded protein responses (Fig. S5A+B). It was detected by POM8 and POM19, which bind to a conformational epitope on the the α1-α2 and β1-α3 regions, respectively (*5*), but not by POM1 (Fig. S3A). The POM1-induced H-latch allosterically altered the β2-α2 loop; similarly, binding of mPrP^2Cys^ to POM5 (recognizing the β2-α2 loop, (*5*)) was impaired (Fig. S3A). Taken together, these suggest that mPrP^2Cys^ adopts a conformation similar to that induced by POM1 (Fig. S4C). We transduced *Prnp*^ZH3/ZH3^ COCS with adeno-associated virus-based vectors (AAV) expressing either PrP^C^ or PrP^2Cys^. Wild-type and mutant proteins showed similarly robust expression levels (Fig. S5C). COCS expressing mPrP^2Cys^ developed spontaneous, dose-dependent neurodegeneration 4 weeks after transduction (Fig. 2A+B, Fig. S6A+B), suggesting that induction of the H-latch suffices to generate toxicity. In agreement with this view, MD simulations showed that human, hereditary PrP mutations responsible for fatal prion diseases favor H-latch formation and altered flexibility in the α2-α3 and β2-α2 loop (Fig. S7).

**Fig. 2.**
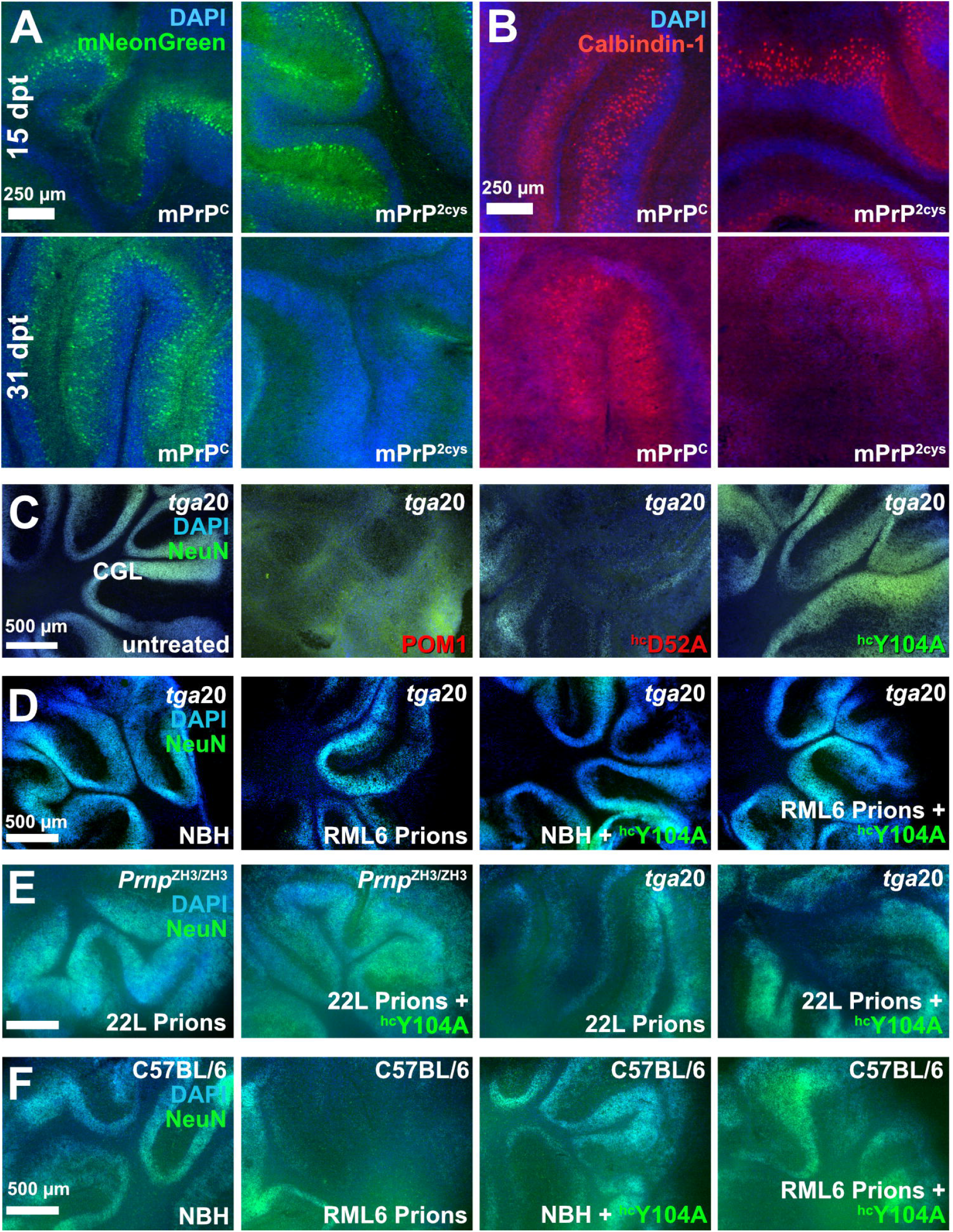
Preventing H-latch formation by pomologs rescues prion-induced neurodegeneration. **(A-B)** *Prnp*^ZH3/ZH3^ COCS transduced with a bi-cistronic AAV expressing mNeonGreen and mPrP^c^ (*left*) or mPrP^2Cys^ (*right*). **(A)** mNeonGreen was visible in all COCS at 15 days post transduction (dpt, *top row*) but disappeared in mPrP^2Cys^ at 31 dpt (*bottom row*). **(B)** Calbindin-1^+^ Purkinje cells were preserved at 15 dpt but became largely undetectable at 31 dpt, as a result of mPrP^2Cys^ toxicity. **(C)** The densely cellular NeuN^+^/DAP^+^ cerebellar granule cell layer (CGL) of *tga20* COCS was preserved by treatment with POM1 mutant ^hc^Y104A (green) but destroyed by POM1 and ^hc^D52A (red). **(D)** CGL degeneration in prion-infected *tga20* COCS but not in COCS exposed to non-infectious brain homogenate (NBH). Treatment of RML6 prion-infected *tga20* COCS with ^hc^Y104A prevented neuronal loss. (E) Rescue of prion-induced toxicity by ^hc^Y104Ain COCS inoculated with 22L prions. (F) Treatment of prion-infected wild-type COCS, expressing wild-type levels of PrP^c^, with ^hc^Y104A prevented CGL degeneration. Quantification of fluorescent micrographs is depicted in Fig. S6.

If POM1 toxicity requires the H-latch, antibody mutants unable to induce it should be innocuous. POM1 immobilizes R208 by salt bridges with its heavy-chain (hc) residue ^hc^D52, whereas ^hc^Y104 contributes to the positioning of H140 (Fig. 1A). To prevent H-latch formation, we thus replaced nine of these residues with alanine. For control, we similarly substituted interface residues predicted to have no impact on R208. Resulting “pomologs” were produced as singlechain variable fragments (scFv), three of which retained high affinity, i.e. K_D_ ≈ 10 nM, for PrP^C^ (Supplementary Table 1, Fig. S8-S9).

As expected, all pomologs were innocuous to *Prnp*^ZH1/ZH1^ COCS that do not express PrP^C^ (*12*) (Fig. S10). ^hc^Y104A reduced H-latch formation according to MD simulations (Fig. 1B, Fig. S2) and exerted no neurotoxicity onto COCS from *tga20* mice overexpressing PrP^C^ (*13*), whereas POM1 and all H-latch inducing mutants (^hc^D52A, ^hc^Y101A and all light-chain pomologs) were neurotoxic (Fig. 2C, Fig. S6C). As with POM1, the toxicity of pomologs required PrP^C^, featured neuronal loss, astrogliosis and elevated levels of microglia markers (Fig. S11A+B), and was ablated by co-administration of the antibody POM2 which targets the flexible tail (FT) of PrP^C^ (Fig. S11C) (*5*). Additionally, ^hc^Y104A inhibited POM1 toxicity (Fig. S12A+B).

POM1 does not induce *de novo* prions (*14*) but triggers similar neurotoxic cascades (*6*), plausibly by replicating the docking of prions to PrP^C^. If so, ^hc^Y104A may prevent the neurotoxicity of both POM1 and prions by competing for their interaction with PrP^C^. Indeed, ^hc^Y104A protected RML6 and 22L prion-inoculated *tg*a20 and C57BL/6 COCS from prion neurodegeneration (Fig. 2D-F and Fig. S6D-F), repressed the vacuolation of chronically prion-infected cells (Fig. S12C and (*1*)) and diminished PrP^Sc^ levels *ex vivo* (Fig. S12D). In contrast to other antiprion antibodies (*15*), ^hc^Y104A did not reduce levels of PrP^C^ (Fig. S12E), corroborating the conjecture that neuroprotection results from interfering with the docking of incoming prions.

Antibody ICSM18 was found to ameliorate prion toxicity *in vivo* (*16*) although dose escalation studies showed conspicuous neuronal loss (*7*). The ICSM18 epitope is close to that of POM1 (*8*), and MD simulations indicated that it facilitates the R208-H140 interaction, albeit less than POM1 (Fig. 1C).

Protective pomolog ^hc^Y104A failed to induce the H-latch compared to toxic ones (Fig. 1C, Fig. S1). MD simulations showed that POM1 rigidified its epitope but increased the flexibility of the α2-α3 and β2-α2 loops (Fig. 1C). Conversely, the conformation of PrP attached to the protective ^hc^Y104A resembled that of free PrP. Consistent with MD simulations, NMR spectra, which are sensitive to local effects and transient populations (*17*), of rmPrP_90-231_ complexed with POM1 revealed long-range alterations in the GD and in the adjacent FT (Fig. 3A). When bound to ^hc^Y104A, instead, rmPrP_90-231_ elicited spectra similar to those of free PrP. Circular-dichroism (CD) spectroscopy showed that the full rmPrP (rmPrP_23-231_)-POM1 complex had more irregular structure content than its free components (Fig. 3B), whereas no difference was observed when POM1 was complexed to partially FT-deficient rmPrP_90-231_. This suggests that POM1 can alter the secondary structure of the FT. We did not observe any changes in the secondary structure of the ^hc^Y104A-bound rmPrP_23-231_ complex. Hence H-latch induction leads to subtle alterations of the structure of both GD and FT, whose presence correlates with toxicity. We performed animal experiments to confirm that i) ^hc^Y104A by itself is not neurotoxic *in vivo*, in contrast to POM1, and ii) it protects from prion-dependent neurodegeneration. When produced as IgG holoantibody, ^hc^Y104A exhibited subnanomolar affinity to full-length, murine, recombinant PrP (rmPrP_23-231_, Fig. S13). We injected POM1 or holo-^hc^Y104A into the hippocampus of C57BL/6 mice. Histology and volumetric diffusion-weighted magnetic resonance imaging showed that POM1 (6 μg) elicited massive neurodegeneration that was repressed by preincubation with recPrP in three-fold molar excess, whereas the same amount of holo-^hc^Y104A did not elicit any tissue damage (Fig. 3C, Fig. S14-S15). A benchmark dose analysis (*7*) yielded an upper safe dose limit of ≥ 12 μg for intracerebrally injected holo-^hc^Y104A (Fig. S16A). Also, the injection of holo-^hc^Y104A (6 μg) into *tga20* mice, which are highly sensitive to POM1 damage, failed to induce any lesions (Fig. S16B-E).

**Fig. 3.**
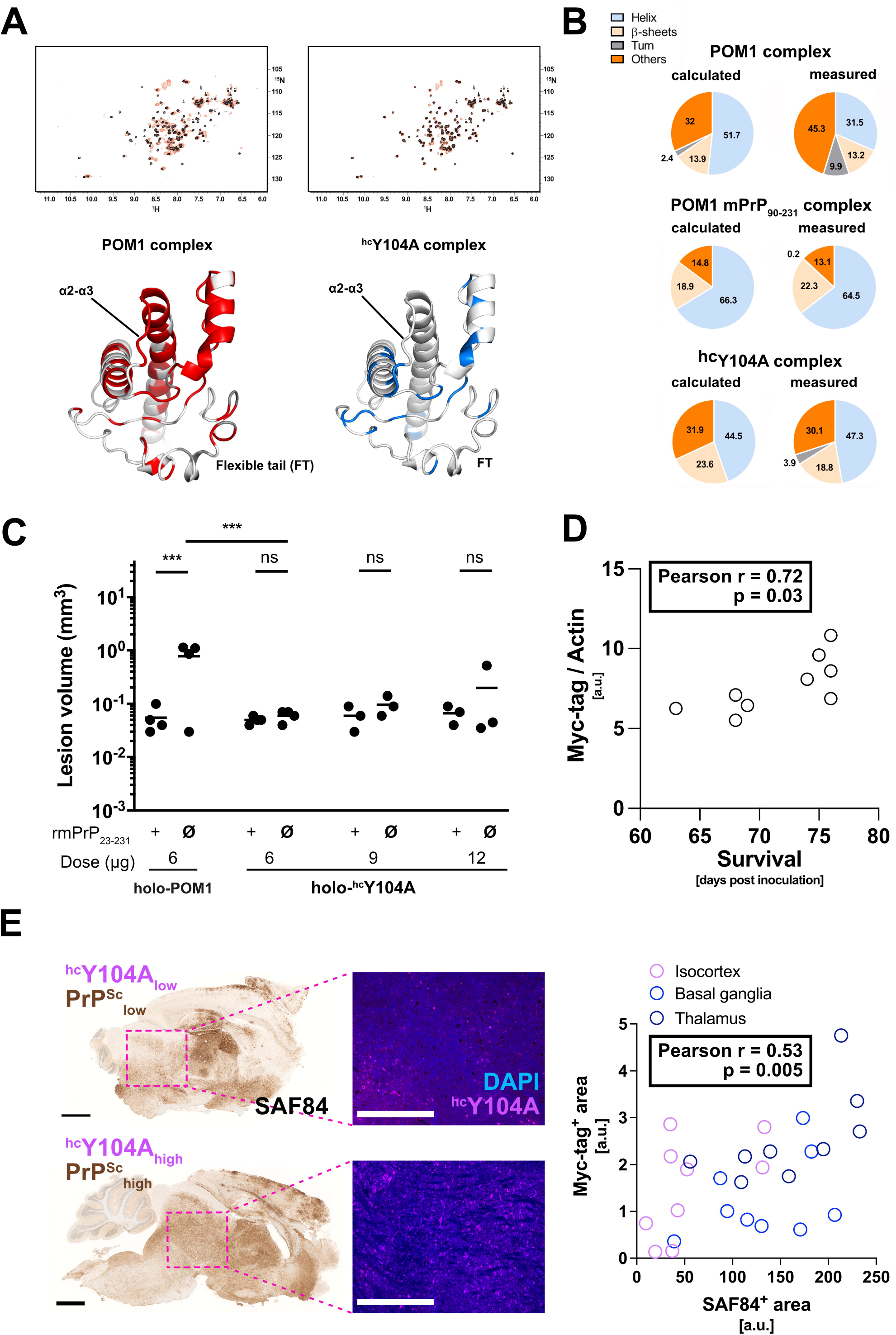
Antibody binding causes allosteric conformational changes in globular domain and flexible tail. **(A)** Comparison between the [^15^N,^1^H]-TROSY spectra of rm-PrP_90-231_ free vs. bound to ^hc^Y104A pomolog. Chemical shift differences, reflecting subtle alterations of the local chemical structure, were visible not only in the epitope but also at distant sites in the GD and FT. Residues affected by antibody binding are in color on PrP^c^ (GD and part of the FT are shown). Differences between toxic and protective antibody are evident in the α2-α3 loop (the Y104A complex is identical to free PrP^c^) and in the FT region closer to the GD. (B) Content of secondary structure estimated from CD spectra of the rmPrP-pomologs complexes. “Calculated” indicates the secondary structure content if rmPrP and pomolog would not change upon binding. POM1 displayed increased content of irregular structure (measured vs. calculated) when complexed with full rmPrP_23-231_ but identical content when complexed with a construct lacking the FT (rmPrP_90-231_). This indicates that the FT changes conformation upon POM1 binding. Conversely, no differences were detected with the protective pomolog ^hc^Y104A. (C) Volumetric quantification of lesions on DWI imaging 24 hours after injection revealed no significant lesion induction by holo-^hc^Y104A. **(D)** Antibody expression levels, as determined by Myc-Tag western blot, showed a positive correlation with survival. (E) Significant correlation of PrP^Sc^ and antibody expression levels levels (right graph, aggregated correlation across all brain regions). Scale bar SAF84: 1 mm. Scale bar ^hc^Y104A-Myc-tag: 500 μm. a.u.: arbitrary units.

We then transduced *tga*20 mice with ^hc^Y104A by intravenous injection of a neurotropic AAV-PHP.B vector. Two weeks after AAV injection, mice were inoculated intracerebrally with 3 x 10^5^ ID_50_ units of RML6 prions. ^hc^Y104A expression levels correlated with both survival times and PrP^Sc^ deposition (Fig. 3D+E) suggesting that ^hc^Y104A acts downstream of prion replication. If the same toxic PrP conformation is induced by both the H-latch and infectious prions, anti-PrP antibodies unable to bind the H-latch conformers could depopulate them by locking PrP^C^ in its innocuous state, thus preventing prion neurotoxicity. Using phage display (Fig. S17) we generated four antigen-binding fragments (Fabs), three of which bound the globular domain of PrP^C^ preferentially over PrP^2Cys^ whereas one bound PrP and PrP^2Cys^ similarly (Fig. 4A; S18). When administered to prion-infected *tga20* COCS, FabA10 and FabD9 decreased prion neurotoxicity whereas FabE2, which binds both PrP^C^ and mPrP^2Cys^, had no beneficial effect (Fig. 4B+C). NMR epitope mapping followed by computational docking and MD (*18*) showed that FabA10 binds to PrP encompassing the H-latch and partially overlapping with the POM1 epitope (Fig. 4D, Fig. S19-S20). MD showed that the H-latch is not stable in the presence of FabA10 even if the simulations were started from a POM1-bound PrP conformation with the R208-H140 H-bond present (Fig. S19).

**Fig. 4.**
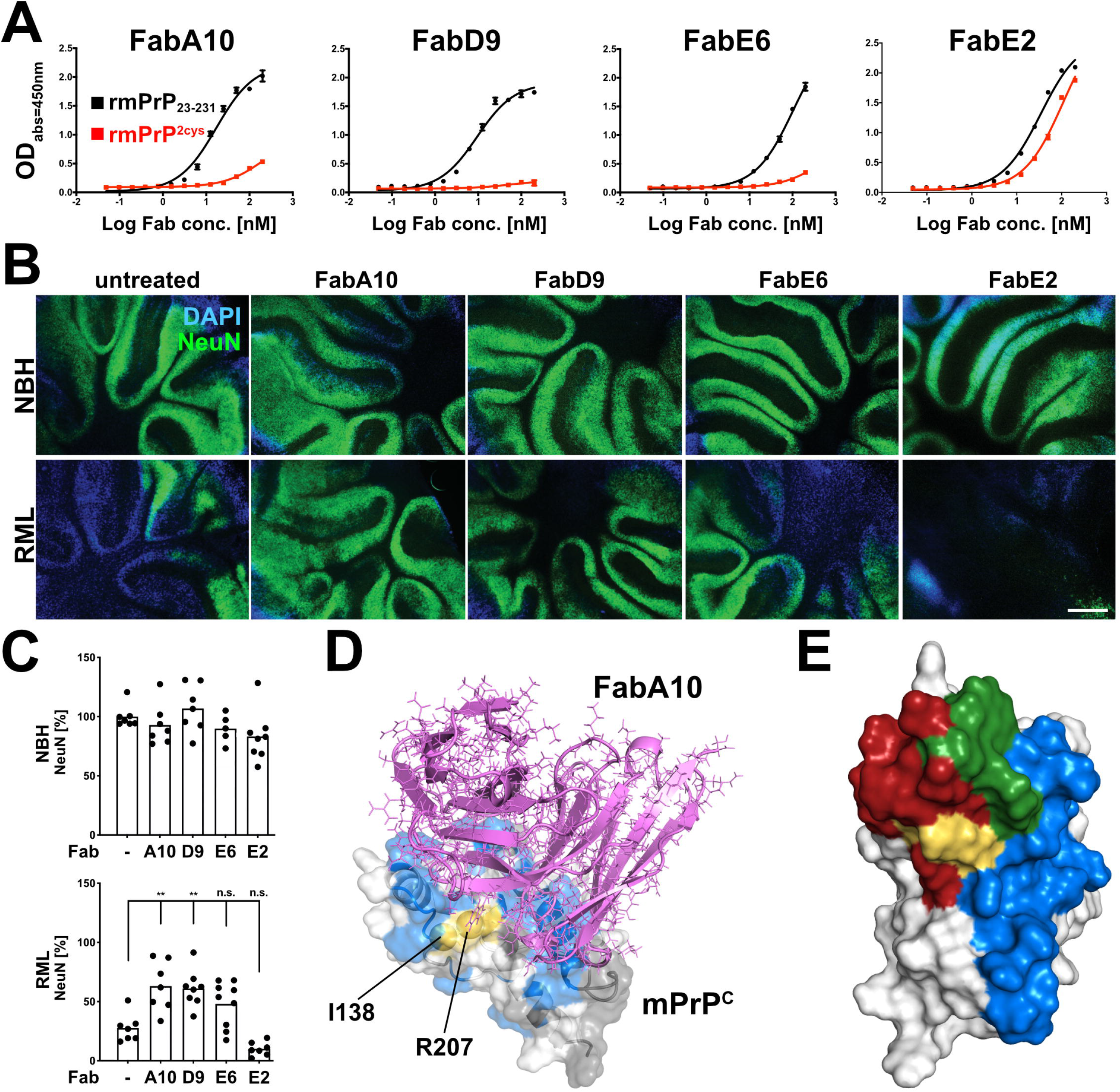
Phage-displayed antibody fragments differentially binding wild-type PrP^c^, but not PrP^2Cys^, confer neuroprotection. **(A)** Preferential binding of the selected Fabs to rmPrP_23-231_ over rmPrP^2Cys^. With the exception of FabE2, the Fabs show higher apparent affinity for rmPrP_23-231_ than rmPrP^2Cys^. (B) FabA10 and FabD9 conferred neuroprotection in prion-infected *tga20* COCS. **(C)** Quantification of NeuN fluorescence intensity from (B), expressed as percentage of untreated (-) NBH. Scale bar = 500 μm. (D) Structure of PrP^c^ (white) in complex with FabA10 (violet) obtained by NMR validated docking and MD. mPrP_90-231_ residues whose NMR signal is affected by FabA10 binding are colored blue; residues with no NMR information grey; residues mutated to Cys are yellow. (E) There is partial overlap (green) between the epitopes of POM1 (red) and FabA10 (blue). The 2 Cys are in yellow. PrP^c^ is depicted in different orientations in D and E. n.s. not significant, ** p < 0.01

In summary, the evidence presented here suggests that H-latch formation is an important driver of prion toxicity. The H-latch was induced by the toxic anti-PrP antibody POM1, PrP mutants unable to form the H-latch conferred resistance to POM1 toxicity, and a PrP mutant mimicking the H-latch was constitutively neurotoxic. Conversely, POM1 mutants retaining its affinity and epitope specificity but abolishing H-latch formation. We observed formation of the H-latch and its structural effects on PrP^C^-GD were not only innocuous but also protective against prion neurotoxicity in vitro and in vivo. The molecular dynamics predictions were confirmed in vivo using both cerebellar slice cultures and mouse models of prion disease. POM1 mutants or other rationally selected Fabs that were unable to induce the H-latch protected from the deleterious effects of prion infection *ex vivo* and *in vivo*. Furthermore, hereditary PrP mutations leading to human prion diseases also favor the H-latch according to MD simulations. These observations suggest that the H-latch is not only involved in the toxicity of anti-PrP antibodies but also in the pathogenesis of prion diseases. Other determinants of prion toxicity besides the H-latch include presence of an intact PrP^C^-FT (*5*) and copper-binding properties of PrP^C^ (*19*) or, possibly, recently described polymorphisms in genes outside of *PRNP* (*20*).

The above findings hold promise for therapeutic interventions. Firstly, the POM1 binding region includes a well-defined pocket created by the α1-α3 helix of PrP^C^, which may be targeted by therapeutic compounds including antibodies, small molecules, cyclic peptides or aptamers. Secondly, ^hc^Y104A halted progression of prion toxicity even when they were already conspicuous, whereas the anti-FT antibody POM2 exerted neuroprotection only when applied directly after prion inoculation (*9*). This suggests that ^hc^Y104A halts prion toxicity upstream of FT engagement (*6, 9*). Thirdly, *tga*20 COCS (which are much more responsive to toxic pomologs than wild-type COCS, and can therefore be regarded as a sensitive sentinel system) tolerated prolonged application of ^hc^Y104A at concentrations around 150 * K_D_. Finally, intracerebrally injected ^hc^Y104A was innocuous, and AAV-transduced ^hc^Y104A extended the life span of prion-infected mice. These findings suggest that blockade of the POM1 epitope by agents that do not induce the H-latch enjoys good *in vivo* tolerability. In view of the reports that PrP^C^ may mediate the toxicity of disparate amyloids (*21*), the relevance of the above findings may extend to proteotoxic diseases beyond spongiform encephalopathies.

## Acknowledgements

We would like to acknowledge Mirka Epskamp, Tina Kottarathil, Manfredi Carta, Melvin Rincon, Rita Moos, Jingjing Guo and Clemence Tournaire for valuable discussions and technical help, as well as Dr. Tiziana Sonati for advising on certain experiments performed by Ms. Tournaire. We are grateful to Giulia Moro for help and discussion. Imaging was performed with equipment maintained by the Center of Microscopy and Image Analysis, University of Zurich. The viral vectors and respective plasmids were produced by the Viral Vector Facility (VVF) of the Neuroscience Center Zurich (Zentrum fu□r Neurowissenschaften Zu□rich, ZNZ). We are grateful to Prof. Ana Paula Valente for useful discussion on protein dynamics.

## Authors contributions

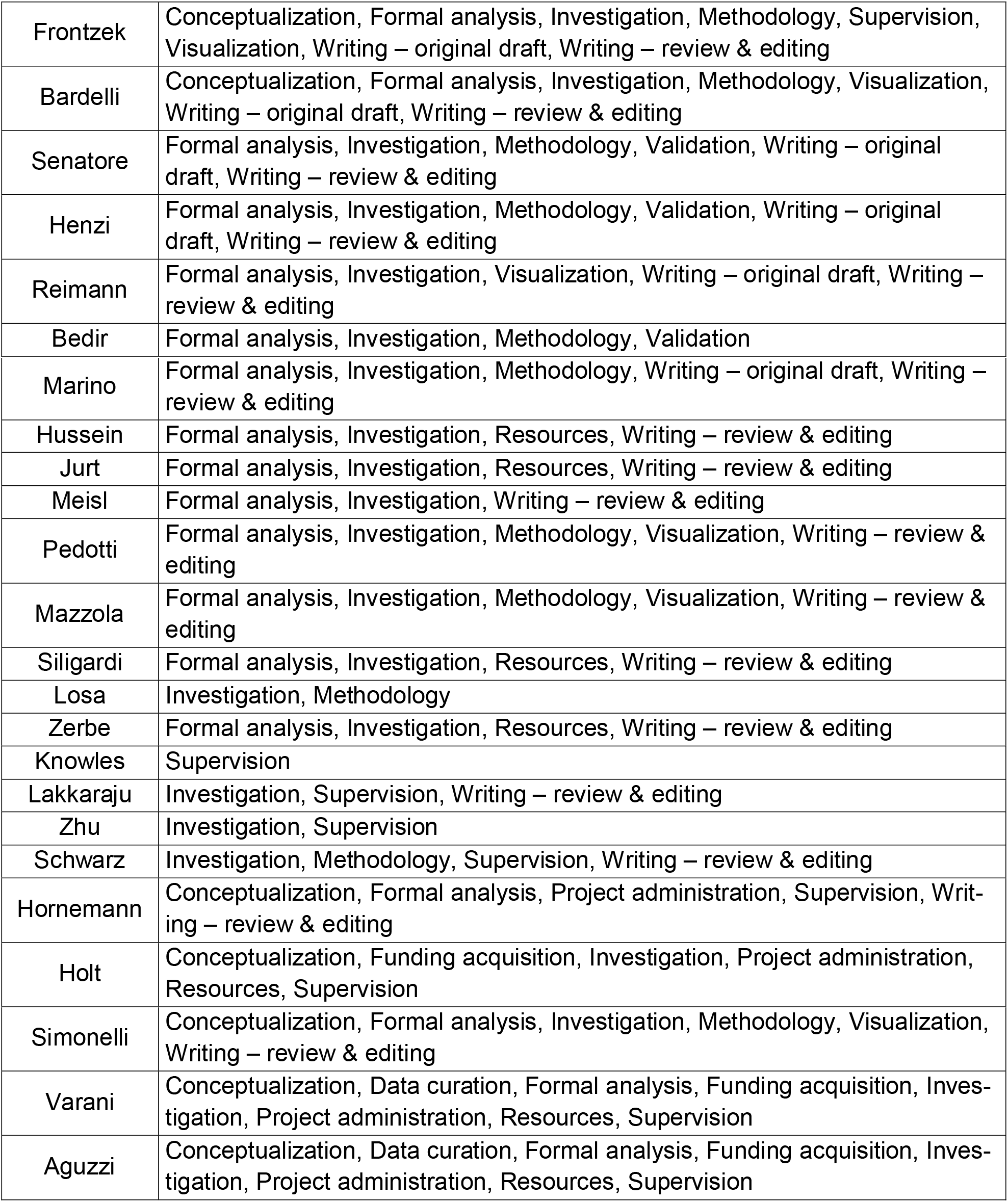

## Funding

KF received unrestricted support by the Theodor und Ida Herzog-Egli-Stiftung and Ono Pharmaceuticals. RR was supported by a Career Development Award from the Stavros Niarchos Foundation. GM is funded by a Ramon Jenkins Research Fellowship at Sidney Sussex College. TK received financial support by the EPSRC, BBSRC, ERC, and the Frances and Augustus Newman Foundation. AA is supported by an Advanced Grant of the European Research Council (ERC, No. 250356). a Distinguished Scientist Award of the Nomis foundation, and grants from the GELU foundation, the Swiss National Foundation (SNF, including a Sinergia grant), and the Swiss Initiative in Systems Biology, SystemsX.ch (PrionX, SynucleiX). LV gratefully acknowledges support from SNF (No. 310030_166445, 157699), Synapsis Foundation_Alzheimer research (ARS) and Lions Club Monteceneri. We would like to thank Diamond Light Source for B23 beamtime allocation (CM-19680). MGH was supported by a grant from the Thierry Latran Foundation (SOD-VIP), FWO (Grant 1513616N), European Research Council (ERC) Proof of Concept Grant 713755 - AD-VIP) and the European Commission (H2020-WIDESPREAD-2018-2020-6; NCBio; 951923). The funders had no role in study design, data collection and analysis, decision to publish, or preparation of the manuscript.

## Competing interests

The authors declare no competing interests.

## Data and materials availability

Auxiliary supplementary data (uncropped gels, FACS gating strategy, gene sequences) are available via FigShare: https://doi.org/10.6084/m9.figshare.11940606. All other data is available in the manuscript or the supplementary materials. All unique biological materials used in the manuscript are readily available from the authors.

## Supplementary Data

Materials and Methods

Supplementary Table S1

Supplementary Figures S1 – S20

## Materials and Methods

### Adeno-associated virus production and in vivo transduction

Single stranded adeno-associated virus (ssAAV) vector backbones with AAV2 inverted terminal repeats (ITRs) were kindly provided by Berhard Schneider (EPFL, Switzerland). Herein, expression of the monomeric NeonGreen (mNG) fluorophore was driven by the human Synapsin I (hSynI) promoter. A P2A sequence (GSGATNFSLLKQAGDVEENPGP) was introduced between mNeonGreen and PrP^C^ for bi-cistronic expression. For mPrP^R207A^ and mPrP^2cys^ expression, a synthetic gene block (gBlock, IDT, full sequence was deposited on FigShare https://doi.org/10.6084/m9.figshare.11940606) was cloned between the BsrGI and HindIII site of the vector replacing the wild-type PrP^C^ sequence. Recombination of plasmids was tested using SmaI digestion prior to virus production. The viral vectors and viral vector plasmids were produced as hybrid AAV2/6 (AAV6 capsid with AAV2 ITRs) by the Viral Vector Facility (VVF) of the Neuroscience Center Zurich (Zentrum fu□r Neurowissenschaften Zu□rich, ZNZ, Switzerland. The identity of the packaged genomes was confirmed by Sanger DNA-sequencing (identity check). Quantification of mNeonGreen-positive cells from confocal images was done using the *Spots* function in Imaris (Bitplane).

Neurotropic AAV variants for scFv antibody expression were constructed from synthetic gene fragments, NheI-IL2-scFv-Myc-EcoRV (produced by Genscript Biotech, New Jersey, USA), that contained ^hc^Y104A sequences preceded by the signal peptide from interleukin-2 (IL-2) (*22*). NheI and EcoRV restriction enzyme digestion was performed on NheI-IL2-scFv-Myc-EcoRV synthetic gene fragments which were then inserted into a ssAAV vector backbone. ScFv expression was under the control of the strong, ubiquitously active CAG promoter. A WPRE sequence (woodchuck hepatitis virus post-transcriptional regulatory element) was also included, downstream of the transgene, to enhance transgene expression. Production, quality control and determination of vector titre was performed by ViGene Biosciences (Rockville, Maryland, USA). Rep2 and CapPHP.B plasmids were provided under a Material Transfer Agreement (MTA). Further details about packaging and purification strategies can be found on the company’s website (http://www.vigenebio.com).

### Allen Mouse Brain Atlas data

Images from *in situ* hybridization for Calbindin 1 and Synapsin 1 expression were taken from the Allen Mouse Brain Atlas (www.brain-map.org). The first dataset retrieved by the R package *allenbrain* (https://github.com/oganm/allenBrain) with the closest atlas image to the center of the region (regionID = 512, settings: planeOfSection = ‘coronal’, probeOrientation = ‘anti-sense’) was downloaded (dataset IDs for calb1 = 71717640, syn1 = 227540). Image credit: Allen Institute.

### Animals and in vivo experiments

We conducted all animal experiments in strict accordance with the Swiss Animal Protection law and dispositions of the Swiss Federal Office of Food Safety and Animal Welfare (BLV). The Animal Welfare Committee of the Canton of Zurich approved all animal protocols and experiments performed in this study (animal permits 123, ZH90/2013, ZH120/16, ZH139/16). Genetically modified mice from the following genotypes were used in this study: Zurich I *Prnp*^o/o^ (denoted as *Prnp*^ZH1/ZH1^) (*12*), Zurich III *Prnp^O/O^* (denoted as *Prnp*^ZH3/ZH3^) (*11*) and *tga20* (*13*).

For *in vivo* transduction with the neurotropic AAV-PHP.B construct, mice received a total volume of 100 μl (1 x 10^11^ total vector genomes) by intravenous injection into the tail vein. 14 days after AAV transduction *Tga*20 mice were inoculated into the left hemisphere with 30 μl of 0.1% RML6 brain homogenate, corresponding to 3 × 10^5^ LD50 (3.6 μg of total brain homogenate, respectively). Brain homogenates were prepared in 0.32 M sucrose in PBS at a concentration of 10% (w/v). Protein analysis of mouse brains is described below.

After fixation with 4% paraformaldehyde for 1 week, tissues were treated with concentrated formic acid for 60 min, fixed again in formalin and eventually embedded in paraffin. HE staining and SAF84 immunohistochemistry were performed as described previously (*23*). For immunohistochemical detection of Myc-tag, tissue was deparaffinized and incubated in citrate buffer (pH 6.0) in a domestic microwave for 20 min. Unspecific reactivity was blocked using blocking buffer (10% goat serum, 1% bovine serum albumin, 0.1% Triton-X100 in PBS) for 1 hour at room temperature. Primary rabbit anti-Myc tag antibody (1:200, ab9106, Abcam, overnight at 4°C) was detected with Alexa Fluor^®^ 594 Rabbit Anti-Goat (IgG) secondary antibody (1:1’000, 1 h at room temperature), diluted in staining buffer (1% bovine serum albumin, 0.1% Triton-X100 in PBS). Tissue was counterstained with DAPI (5 μg/ml, 15 min at room temperature).

### Cell lines

CAD5 is a subclone of the central nervous system catecholaminergic cell line CAD showing particular susceptibility to prion infection (*14*). Generation of the CAD5 *Prnp*^-/-^ clone #C12 was described before, as was overexpression of murine, full-length PrP^C^ in CAD5 *Prnp*^-/-^ by cloning the open reading frame of *Prnp* into the pcDNA3.1(+) vector, *Prnp* expression was driven by a constitutively expressed cytomegalovirus promoter (yielding pcDNA3.1(+)-*Prnp*) as described earlier (*9*). For stable expression of mPrP^2cys^, pcDNA3.1(+)-*Prnp* vector was modified using Quikchange II Site-Directed Mutagenesis Kit (Agilent) according to the manufacturer’s guidelines. We first introduced a mutation leading to R207C (primers (5’ -> 3’): mutagenesis FW: GTG-AAG-ATG-ATG-GAG-TGC-GTG-GTG-GAG-CAG-A, REV: TCT-GCT-CCA-CCA-CGC-ACT-CCA-TCA-TCT-TCA-C) which was then followed by mutation of I138C (mutagenesis FW: AGT-CGT-TGC-CAA-AAT-GGC-ACA-TGG-GCC-TGC-TCA-TGG, REV: CCA-TGA-GCA-GGC-CCA-TGT-GCC-ATT-TTG-GCA-ACG-ACT). For stable expression of mPrP^R207A^, pcDNA3.1(+)-*Prnp* was mutated correspondingly (mutagenesis FW: TGT-GAA-GAT-GAT-GGA-GGC-CGT-GGT-GGA-GCA-GAT-G, REV: TCT-GCT-CCA-CCA-CGC-ACT-CCA-TCA-TCT-TCA-C).

### Cell vacuolation assay

Mouse hypothalamic Gt1 neuronal cells were grown in Dulbecco’s modified eagle medium (DMEM) in the presence of 10% foetal bovine serum (FBS), 1% penicillin-streptomycin and 1% Glutamax (all obtained from Invitrogen). For prion infection of the cells, Gt1 cells growing in DMEM medium were incubated with either Rocky mountain laboratory strain of prion (RML6) prions (0.1%) or non-infectious brain homogenate (NBH; 0.1%) for 3 days in one well of a 6 well plate. This was followed by splitting the cells at 1:3 ratio every three days for at least 10 passages. The presence of infectivity in the cells was monitored by the presence of proteinase K (PK) resistant PrP, as described below. At 70 dpi, the cells started developing vacuoles which were visualized by phase contrast microscopy. Antibody treatment with ^hc^Y104A was administered on 70-75 dpi at a concentration of 180 nM.

### Cerebellar organotypic slice cultures (COCS)

Mice from C57BL/6, *tga*20, *Prnp*^ZH1/ZH1^and *Prnp*^ZH3/ZH3^ strains were used for preparation of COCS as described (*10*). Herein, 350 μm thick COCS were prepared from 9-12 day old pups. Prion infection of COCS was done as free-floating sections with 100 μg per 10 slices of RML6 (= passage 6 of the Rocky Mountain Laboratory strain mouse-adapted scrapie prions) or 22L (mouse-adapted scrapie prions) brain homogenate from terminally sick prion-infected mice. Brain homogenate from CD1-inoculated mice was used as non-infectious brain homogenate (NBH). Sections were incubated with brain homogenates diluted in physiological Grey’s balanced salt solution for 1 h at 4°C, then washed and 5-10 slices were placed on a 6-well PTFE membrane insert. Analogously, for AAV experiments, COCS were incubated with AAV at a final concentration of 5.2 x 10^10^ total vector genomes diluted in physiological Grey’s balanced salt solution for 1 h at 4°C, then washed and placed on PTFE membrane inserts. Antibody treatments were given with every medium change at the designated time periods. In naïve slices, antibody treatments were initiated after a recovery period of 10-14 days.

For testing of innocuity of pomologs (Fig. 2C, Fig. S7C, Fig. S10), POM1 and pomolog antibodies were added at 400 nM for 14 days. Figures S7C and S10 represent aggregated data from multiple experiments with COCS from mice of identical genotype and age, compounds were administered at identical timepoints and dosage. When added to RML-infected *tga20* COCS (Fig. 2D, Fig. S7D), ^hc^Y57A was added at 20 dpi, ^hc^Y104A was added at 21 dpi, both antibodies were given at 400 nM until 45 dpi. Antibody treatment with ^hc^Y57A and ^hc^Y104A of RML-infected *tga20* COCS used for determination of PrP^Sc^ was initiated and stopped at 21 dpi and 45 dpi, respectively (Fig. S13). ^hc^D55A was added to RML-infected *tga*20 COCS at either 1 (800 nM, Fig. S12D) or 21 dpi (400 nM, Fig. 2D, Fig. S7D). When added to C57BL/6 COCS (Fig. 2E, Fig. S7E), hc^Y104A^ was added from 1 dpi at 400 nM until 45 dpi. In 22L inoculated COCS, hcY104A was administered at 21 dpi and slices were harvested at 44 dpi. Phage-derived Fabs were added to RML-infected COCS (Fig. 4B+C) from 1 dpi until 45 dpi at 550 nM.

### Enzyme-linked immunosorbent assay (ELISA)

PrP^C^ levels were measured by ELISA using monoclonal anti-PrP^C^ antibody pairs POM19/POM3 or POM3/POM2 (all as holo-antibodies) as described previously (*24*). 384-well SpectraPlates (Perkin Elmer) were coated with 400 ng mL^-1^ POM19 (POM3) in PBS at 4°C overnight. Plates were washed three times in 0.1% PBS-Tween 20 (PBS-T) and blocked with 80 μl per well of 5% skim milk in 0.1% PBS-T for 1.5 h at room temperature. Blocking buffer was discarded and samples and controls were added dissolved in 1% skim milk in 0.1% PBS-T for 1 h at 37°C. 2-fold dilutions of rmPrP_23-231_, starting at a dilution of 100 ng/ml in 1% skim milk in 0.1% PBS-T were used as calibration curve. Biotinylated POM3 (POM2) was used to detect PrP^C^ (200 ng/ml in 1% skim milk in 0.1% PBS-T), biotinylated antibody was detected with Streptavidin-HRP (1:1’000 in 1% skim milk in 0.1% PBS-T, BD Biosciences). Chromogenic reaction and reading of plates were performed as described in (*24*). Unknown PrP^C^ concentrations were interpolated from the linear range of the calibration curve using linear regression (GraphPad Prism, GraphPad Software).

### ELISA screening of phage display

Single colonies were picked and cultured in 384 well plate (Nunc) in 2YT/Ampicillin/1% glucose medium over night at 37°C, 80% humidity, 500 rpm. These precultures were used to prepare glycerol stock master plates. Expression plates were prepared from the master plates by inoculating corresponding wells with 2YT/Carbenicillin/0.1% glucose medium, followed by induction with 1 mM IPTG. After 4 h at 37°C, 80% humidity, cultures were lysed for 1.5 h at 400 rpm, 22°C in borate buffered saline pH 8.2 containing EDTA-free protease inhibitor cocktail, 2.5 mg/ml lysozyme and 40 U/ml benzonase. Fab-containing bacteria lysate was blocked with Superblock and used for ELISA screening, here, the reactivity to four different antigens was assessed in parallel. The following antigens were coated on separate 384-well ELISA plates: anti-Fd antibody (The Binding Site GmbH) 1:1000 in PBS, to check the expression level of each Fab clone in bacteria; rmPrP_23-231_ at 87 nM in PBS, to identify candidate PrP^C^ binders; mPrP^2cys^ at 87 nM in PBS, to check for cross reactivity with mPrP^2cys^; neutravidin at 87 nM as a control for specificity. Antigen-coated ELISA plates were washed twice with PBS-T and blocked with Superblock for 2 h. Fab containing bacteria lysates from the expression plate were transferred to corresponding wells of the ELISA plates. After 2 h incubation, ELISA plates were washed three times with PBS-T and anti-human F(ab’)2-alkaline phosphatase conjugated antibody (1:5000 in PBS-T) was added. After 1 h incubation at RT, followed by three washings with PBS-T, pNPP substrate was added and, after 5 min incubation, the ELISA signal was measured at 405 nm. Fabs from bacteria lysates producing an ELISA signal 5 times higher than the technical background, which was calculated as the average of the coated well containing un-inoculated medium, and negative for neutravidin were considered as PrP^C^ binder candidates. For hit selection, we only considered anti-PrP^C^ Fabs whose ELISA signal for rmPrP_23-231_ was at least 2 times higher than for mPrP^2cys^. All the identified hits were checked in a confirmatory ELISA screening. Bacteria cultures of the selected clones were used for DNA minipreps followed by Sanger sequencing using the following sequencing primers: HuCAL_VH (5’-GATAAGCATGCGTAGGAGAAA-3’) and M13Rev (5’-CAGGAAACAGCTATGAC-3’).

### Expression and purification of selected anti-PrP Fabs

Chemical competent BL21(D3) cells (Invitrogen) were transformed with selected pPE2-Fab plasmids and grown on LBagar/Kanamycin/1% glucose plates. A single colony was inoculated into 20 ml of 2xYT/Kanamycin/1% glucose pre-culture medium and incubated for at least 4 h at 37°C, 220 rpm. One litre of 2YT-medium containing Kanamycin/0.1% glucose was inoculated with 20 ml pre-culture and Fab expression was induced by 0.75 mM IPTG followed by incubation over night at 25°C, 180 rpm. The overnight culture was centrifuged at 4000 x g at 4°C for 30 min and the pellet was frozen at −20°C. For Fab purification, thawed pellet was resuspended into 20 ml lysis buffer (0.025 M Tris pH 8; 0.5 M NaCl; 2 mM MgCl_2_; 100 U/ml Benzonase (Merck); 0.25 mg/ml lysozyme (Roche), EDTA-free protease inhibitor (Roche)) and incubated for 1 h at room temperature at 50 rpm. Lysate was centrifuged at 16000 x g at 4°C for 30 min and supernatant was filtrated through 0.22 μM Millipore Express^®^Plus Membrane. Fab purification was achieved via the His6-Tag of the heavy chain by IMAC. Briefly, after equilibration of Ni-NTA column with running buffer (20 mM Na-phosphate buffer, 500 mM NaCl, 10 mM Imidazole, pH 7.4), the bacteria lysate was loaded and washed with washing buffer (20 mM Na-phosphate buffer, 500 mM NaCl, 20 mM Imidazole, pH 7.4). The Fab was eluted with elution buffer (20 mM Na-phosphate buffer, 500 mM NaCl, 250 mM Imidazole, pH 7.4). Buffer exchange was performed using PD-10 columns, Sephadex G-25M (Sigma) whereby the Fab was eluted with PBS.

### Förster Resonance Energy Transfer (FRET)

Europium (Eu^3+^) donor fluorophore was coupled to POM1 (yielding POM1-Eu^3+^) and allophycocyanin (APC) acceptor fluorophores was coupled to holoantibody POM3 (yielding holo-POM3-APC) as previously described (*25*). Full-length, recombinant mouse prion protein (rmPrP_23-231_) was added at a final concentration of 1.75 nM followed by addition of holo-POM3-APC at a final concentration of 5 nM and subsequent incubation at 37°C for 30 minutes whilst constantly shaking at 400 rpm. Pomologs were then added in serial dilutions from 0 to 3 nM and again incubated at 37°C for 60 minutes whilst constantly shaking at 400 rpm, followed by addition of POM1-Eu^3+^ at a final concentration of 2.5 nM). Net FRET was calculated as described previously (*25*).

### Determination of binding constants from FRET

The dependence of the FRET signal on POM1 concentration was modelled by a simple competitive binding model. The binding constant of the FRET labelled POM1-Eu^3+^ was defined as

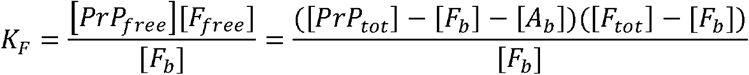

where square brackets denote concentration, *F_tot_, F_free_* and *F_b_* denote total, free and bound POM1-Eu^3+^, *PrP_tot_* and *PrP_free_* denote the total and free PrP, *A_tot_*, *A_free_* and *A_b_* denote total, free and bound scFvs and *K_F_* is the binding constant of the POM1-Eu^3+^. The righthand equality is obtained by imposing conservation of mass. An equivalent equation defines the binding constant of the scFvs

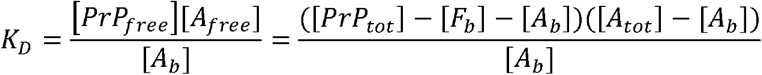

This system of equations is solved to give *F_b_* as a function of *A_tot_*. To relate the concentration of bound POM1-Eu^3+^, *F_b_*, to the FRET measurements this equation was rescaled to 100 for the fully bound and 10 for fully unbound limit. An additional complication in interpreting the experimental data stems from the fact that a FRET signal will only appear if both a POM1-Eu^3+^and holo-POM3-APC are bound to the same PrP. We assume that the binding of POM1 and POM3 is independent, so we can approximate the concentration of PrP bound to a holo-POM3-APC as the effective PrP concentration, *PrP_tot_* in the above equations. The binding constant of holo-POM3-APC was determined to be 0.23 nM, giving an effective concentration of PrP of 1.64 nM (compared to the total PrP concentration of 1.75 nM). To verify the robustness of these results, we also fitted the data assuming a much weaker binding of holo-POM3-APC with a binding constant of 1 nM. The obtained K_D_s of the single-chain fragments were within error of the ones determined with a holo-POM3-APC binding constant of 0.23 nM.

### Immunohistochemical stainings and analysis of immunofluorescence

COCS were washed twice in PBS and fixed in 4% paraformaldehyde for at least 2 days at 4°C and were washed again twice in PBS prior to blocking of unspecific binding by incubation in blocking buffer (0.05% Triton X-100 vol/vol, 0.3% goat serum vol/vol in PBS) for 1 h at room temperature. For visualization of neuronal nuclei, the monoclonal mouse anti-NeuN antibody conjugated with Alexa-488 (clone A60, Life Technologies) was dissolved at a concentration of 1.6 μg mL^-1^ into blocking buffer and incubated for 3 days at 4°C. Further primary antibodies used were recombinant anti-calbindin antibody (1 μg mL^-1^, ab108404, Abcam), anti-glial fibrillary acidic protein (1:500, Z0334, DAKO) and anti-F4/80 (1 μg mL^-1^, MCAP497G, Serotec). Unconjugated antibodies were dissolved in blocking buffer and incubated for 3 days at 4°C. After three washes with PBS for 30 min, COCS were incubated for 3 days at 4°C with secondary antibodies Alexa594-conjugated goat anti-rabbit IgG (Life Technologies) or Alexa647-conjugated goat anti-rat IgG (Life Technologies) at a dilution of 1:1’000 in blocking buffer. Slices were then washed with PBS for 15 min and incubated in DAPI (1 μg mL^-1^) in PBS at room temperature for 30 min to visualize cell nuclei. Two subsequent washes in PBS were performed and COCS were mounted with fluorescence mounting medium (DAKO) on glass slides. NeuN, GFAP, F4/80 and Calbindin morphometry was performed by image acquisition on a fluorescence microscope (BX-61, Olympus), analysis was performed using gray-level auto thresholding function in ImageJ (www.fiji.sc). Cell numbers in supplementary figure 4E were determined using “Spots” function in Imaris (Oxford Instruments). Morphometric quantification was done on unprocessed images with identical exposure times and image thresholds between compared groups. Representative fluorescent micrographs in main and supplementary figures have been processed (linear adjustment of brightness and contrast) for better interpretability.

For immunohistochemistry of CAD5 cells, cells were seeded on 18-well μ-slides (Ibidi) and fixed with 4% paraformaldehyde for 5 minutes at room temperature. Unspecific reactions were blocked using 3% goat serum in PBS for 1 hour at room temperature. Mouse monoclonal anti-PrP^C^ antibodies POM1, POM5, POM8 and POM19 (all holo-antibodies) were established before (*24*), POM antibodies were incubated at 4 μg mL^-1^ in 3% goat serum in PBS at 4°C followed by three washes in PBS. Antibodies were detected using Alexa488-conjugated goat anti-mouse IgG at 1:250 dilution, followed by nuclear counterstain with DAPI (1 μg mL^-1^ in PBS) for 5 minutes at room temperature. Image analysis was performed using SP5 confocal microscope (Leica) with identical exposure times across different experimental groups.

### In vitro toxicity assessment

Quantification of POM1 toxicity on CAD5 *Prnp*^-/-^ stably transfected with mPrP^C^, mPrP^C^_R207A_ or empty control vector as described above was measured as percentage of PI positive cells using Flow Cytometry as described before (*9*).

CAD5 cells were cultured with 20mL Corning Basal Cell Culture Liquid Media-DMEM and Ham’s F-12, 50/50 Mix supplemented with 10% FBS, Gibco MEM Non-Essential Amino Acids Solution 1X, Gibco GlutaMAX Supplement 1X and 0.5mg/mL of Geneticin in T75 Flasks ThermoFisher at 37°C 5% CO2. 16 hours before treatment, cells were split into 96wells plates at 25000 cells/well in 100μL.

POM1 alone was prepared at 5 μM final concentration, in 20 mM HEPES pH 7.2 and 150 mM NaCl. 100 μL of each sample, including buffer control, were added to CAD5 cells, in duplicates.

After 48 hours, cells were washed two times with 100μL MACS buffer (PBS + 1% FBS + 2 mM EDTA) and resuspended in 100 μL MACS buffer. 30’’ before FACS measurements PI (1 μg/mL) was added to cells. Measurements were performed using BD LSRFORTESSA. Percentage of PI positive cells were plotted in columns as mean with SD. Gating strategy is depicted in auxiliary supplementary figures https://doi.org/10.6084/m9.figshare.11940606.

### In vivo toxicity assessment

The in vivo toxicity assessment was performed as previously described (*7*). In brief, mice where i.c. injected by the use of a motorized stereotaxic frame (Neurostar) at the following Bregma coordinates (AP-2 mm, ML ±1.7 mm, DV 2.2 mm, angle in ML/DV plane 15°). Antibodies (2 μl) were injected at a flow rate of 0.5 μl/min. After termination of the injection, the needle was left in place for 3 min.

Mice were placed 24 hours after stereotactic injection on a bed equipped with a mouse whole-body radio frequency transmitter coil and a mouse head surface-coil receiver and then transferred into the 4.7 Bruker Pharma scan. For DWI, routine gradient echo sequences with the following parameters were used: TR: 300 ms TE: 28 ms, flip angle: 90 deg, average: 1, Matrix: 350 x 350, Field of View: 3 x 3 cm, acquisition time: 17 min, voxel size: 87×87 μm^3^, slice thickness: 700 μm, Isodistance: 1400 μm^3^ and b values: 13, 816 s/mm^2^. Finally, mice were euthanized after 49 hours and the brains were fixed in 4% formalin. Coronal section from the posterior cortex were paraffin embedded (4mm) and 2 μm coronal step sections (standard every 100 μm) were cut, deparaffinized and routinely stained with hematoxylin and eosin.

Dose response analysis and the benchmark dose relation were calculated with benchmark dose software (BMDS) 2.4 (United States Environmental Protection Agency).

### Molecular Dynamics

Experimental structures were used as basis for molecular dynamics (MD) simulations when available (scPOM1:mPrP complex, PDB 4H88; free mPrP 1XYX). The structure of full length mPrP, mPrP_□90-231_ and the pomologs was predicted by homology modelling I-Tasser webserver (*26*), based on the experimental structure of the PrP globular domain (aa 120-231) and further validated with MD.

In all simulations the system was initially set up and equilibrated through standard MD protocols: proteins were centered in a triclinic box, 0.2 nm from the edge, filled with SPCE water model and 0.15M Na^+^Cl^-^ ions using the AMBER99SB-ILDN protein force field; energy minimization followed. Temperature (298K) and pressure (1 Bar) equilibration steps of 100ps each were performed. 3 independent replicates of 500ns MD simulations were run with the above-mentioned force field for each protein or complex. MD trajectory files were analyzed after removal of Periodic Boundary Conditions. The overall stability of each simulated complex was verified by root mean square deviation, radius of gyration and visual analysis according to standard procedures. Structural clusters, atomic interactions and Root Mean Square Fluctuation (RMSF) were analyzed using GROMACS (*27*) and standard structural biology tools. RMSF provides a qualitative indication of residue level flexibility, as shown in Fig 1C.

The presence of H-bonds or other interactions between GD residues was initially estimated by visual analysis and then by distance between appropriate chemical groups during the simulation time.

### NMR

Spectra were recorded on a Bruker Avance 600 MHz NMR spectrometer at 298 K, pH 7 in 50mM sodium phosphate buffer at a concentration of 300 □M. In mapping experiments mPrP was uniformly labelled with ^15^N (99%) and ^2^H (approx. 70%), antibodies were unlabelled. PrP and Ab samples were freshly prepared and extensively dialyzed against the same buffer prior to complex formation. The same procedure was followed for CD measurements. Chemical shift assignment was based on published data (BMRB entry 16071)(*28*). Briefly, overlay of [^15^N,^1^H]-TROSY spectra of free or bound mPrP_90-231_ allowed identification of PrP residues for which the associated NMR signal changed upon complex formation, indicating alterations in their local chemical environment (*17*).

### Phage display

A synthetic human Fab phagemid library (Novartis Institutes for BioMedical Research) was used for phage display. First, two rounds of selection against PrP^C^ were performed by coating 96-well Maxisorp plates (Nunc) with decreasing amount of rmPrP_23-231_ (1 μM and 0.5 μM respectively, in PBS), overnight at 4°C. PrP-coated plates were washed three times with PBS-T and blocked with Superblock for 2 h. Input of 4 x 10^11^ phages in 300 μl of PBS was used for the first round of panning. After 2 h blocking with Chemiblocker (Millipore), the phages were incubated with PrP-coated wells for 2 h at room temperature. The non-binding phages were then removed by extensive washing with PBS-T while rmPrP_23-231_ bound phages were eluted with 0.1 M Glycine/HCl, pH 2.0 for 10 min at room temperature, the pH was then neutralized by 1 M Tris pH 8.0. Eluted phages were used to infect exponentially growing amber suppressor TG1 cells (Lubio Science). Infected bacteria were cultured in 2YT/Carbenicillin/1% glucose medium overnight at 37°C, 200 rpm and superinfected with VCSM13 helper phages. The production of phage particles was then induced by culturing the superinfected bacteria in 2YT/Carbenicillin/Kanamycin medium containing 0.25 mM isopropyl β-D-1-thiogalactopyranoside (IPTG), overnight at 22°C, 180 rpm. Supernatant containing phages from the overnight culture was used for the second panning round. Output phages from the second round were purified by PEG/NaCl precipitation, titrated, and used in the following third rounds to enrich phage-displayed Fabs that bound preferentially mPrP^C^ over mPrP^2cys^.

Two strategies were used: depletion of binders to recombinant mPrP^2cys^ by subtraction in solid phase and depletion of mPrP^2cys^ binders by competition with rhPrP^C^_23-230_-AviTag™ in liquid phase. In the former setting, purified phages were first exposed to 0.75 μM mPrP^2cys^ (3-fold molar excess compared to rmPrP_90-231_ or rmPrP_121-231_), and then the unbound phages were selected for rmPrP_90-231_ or rmPrP_121-231_ binders. Alternatively, purified phages were first adsorbed on neutravidin-coated wells to remove the neutravidin binders and then exposed to 0.25 μM rhPrP^C^_23-230_-AviTag^TM^ in solution in the presence of 0.75 μM (3-fold molar excess) of mPrP^2cys^. The phage-displayed Fabs binding to rhPrP^C^_23-230_-AviTag^TM^ were captured on neutravidin-coated wells and eluted as described above. For both strategies, a fourth panning round was performed using 0.3 μM mPrP^2cys^ for depletion and 0.1 μM rmPrP_121-231_ (coated on the plate) or rhPrP^C^_23-230_-AviTag^TM^ (in solution) for positive selection. At the fourth round of selection, DNA minipreps were prepared from the panning output pools by QIAprep Spin Miniprep kit (Qiagen) and the whole anti-PrP Fab enriched library was subcloned in expression vector pPE2 (kindly provided by Novartis, Switzerland). DNA was then used to transform electrocompetent non-amber suppressor MC1061 bacteria (Lubio Science) to produce soluble Fabs and perform ELISA screening.

### Production of recombinant proteins and antibodies

Bacterial production of recombinant, full-length mouse PrP_23-231_, recombinant fragments of human and mouse PrP and recombinant, biotinylated human PrP^C^-AviTag^TM^ (rhPrP^C^_23-230_-AviTag™) was done as previously described (*29–31*). Production of scFv and IgG POM1 anti-bodies used in this manuscript was performed as described before (*24*). Production of holo-^hc^Y104A was undertaken as follows: POM1 IgG_1_ heavy chain containing a Y104A mutation and POM1 kappa light chain were ordered as a bicistronic synthetic DNA block (gBlock, IDT) separated by a P2A site. The synthetic gene block (gBlock, IDT, see full sequence on FigShare https://doi.org/10.6084/m9.figshare.11940606) was then cloned into pcDNA™3.4-TOPO^®^ vector (Thermo Fisher Scientific) and recombinant expression was undertaken using the FreeStyle™ MAX 293 Expression System (Thermo Fisher Scientific) according to the manufacturer’s guidelines. Glucose levels were kept constant over 25 mM. Seven days after cell transfection, medium supernatant was harvested, centrifuged and filtered. A Protein-G column (GE Healthcare) was used for affinity purification of antibodies, followed by elution with glycine buffer (pH = 2.6) and subsequent dialysis against PBS (pH = 7.2-7.4). Purity was determined by SDS-PAGE and protein concentrations were determined using Pierce BCA Protein Assay Kit (Thermo Fisher Scientific). For generation of POM1 mutants, we performed site-directed mutagenesis on POM1 pET-22b(+) (Novagen) expression plasmid (*5*) according to the manu-facturer’s guidelines (primers (5’ -> 3’): hcW33A: FW:

CATTCACTGACTACGCGATGCACTGGGTGAAGC, REV:

GCTTCACCCAGTGCATCGCGTAGTCAGTGAATG. hcD52A: FW:

GAGTGGATCGGATCGATTGCGCCTTCTGATAG, REV:

CTATCAGAAGGCGCAATCGATCCGATCCACTC. hcD55A: FW

GGATCGATTGATCCTTCTGCGAGTTATACTAGTCAC, REVGTGACTAGTATAACTCGCAGAAGGATCAATCGATCC. hcY57A: FW:

CCTTCTGATAGTGCGACTAGTCACAATGAAAAGTTCAAGG, REV:

CCTTGAACTTTTCATTGTGACTAGTCGCACTATCAGAAGG. lcS32A: FW:

CCAGTCAGAACATTGGCACAGCGATACACTGGTATCAGCAAAG, REV:

CTTTGCTGATACCAGTGTATCGCTGTGCCAATGTTCTGACTGG. lcY50A: FW:

CTCCAAGGCTTATCATAAAGGCGGCTTCTGAGTCTATCTCTGG, REV:

CCAGAGATAGACTCAGAAGCCGCCTTTATGATAAGCCTTGGAG. lcS91A: FW:

CAGATTATTACTGTCAACAAGCTAATACCTGGCCGTACACGTT, REV:

AACGTGTACGGCCAGGTATTAGCTTGTTGACAGTAATAATCTG. lcW94A: FW:

GTCAACAAAGTAATACCGCGCCGTACACGTTCGGAGG, REV:

CCTCCGAACGTGTACGGCGCGGTATTACTTTGTTGAC. lcY96A: FW:

TAATACCTGGCCGGCCACGTTCGGAGGGG, REV:

CCCCTCCGAACGTGGCCGGCCAGGTATTA. hcY101A: FW:

CTGTTCAAGATCCGGCGCCGGATATTATGCTATGGAG, REV:

CTCCATAGCATAATATCCGGCGCCGGATCTTGAACAG. hcY104A: FW:

CCGGCTACGGATATGCTGCTATGGAGTACTGGG, REV:

CCCAGTACTCCATAGCAGCATATCCGTAGCCGG), followed by subsequent expression and purification as was described for holo-POM1.

### Protein analysis

COCS were washed twice in PBS and scraped off the PTFE membranes with PBS. Homogenization was performed with a TissueLyser LT (Qiagen) for 2 minutes at 50 Hz. A bicinchoninic acid assay (Pierce™ BCA protein assay kit, Thermo Fisher Scientific) was used to determine protein concentrations. PrP^Sc^ levels were determined through digestion of 20 μg of COCS homogenates with 25 μg mL^-1^ of proteinase K (PK, Roche) at a final volume of 20 μL in PBS for 30 minutes at 37°C. PK was deactivated by addition of sodium dodecyl sulfate-containing NuPAGE LDS sample buffer (Thermo Fisher Scientific) and boiling of samples at 95°C for 5 minutes. Equal sample volumes were loaded on Nu-PAGE Bis/Tris precast gels (Life Technologies) and PrP^C^ / PrP^Sc^ was detected by Western blot using the monoclonal anti-PrP antibodies POM1, POM2, or POM19 at 0.4 μg mL^-1^ (all holo-antibodies) as established elsewhere (*6*). Further primary antibodies used for western blots in this manuscript are as follows: monomeric NeonGreen (1:1’000, Chromotek), phospho-eIF2α (1:1’000, clone #D9G8, Cell Signaling Technologies), eIF2α (1:1’000, clone #D7D3, Cell Signaling Technologies), pan-actin (1:10’000, clone #C4, Millipore), GFAP (1:1’000, clone #D1F4Q, Cell Signaling Technologies), Iba1 (1:500, catalogue # 019-19741, Wako), NeuN (0.5 μg/ml, catalogue # ABN78, Merck Millipore), Myc-tag (1:500, catalogue # ab9106, Abcam). After incubation of primary antibody at 4°C overnight, membranes were washed and detected with a goat polyclonal anti-mouse (1:10’000, 115-035-062, Jackson ImmunoResearch) or goat polyclonal anti-rabbit (1:10’000, 111-035-045, Jackson ImmunoResearch) for 1 hour at room temperature. For PNGaseF digestion, 20 μg of samples were processed using a commercially available kit (New England Biolabs), PrP^C^ detection was performed using the monoclonal anti-PrP^C^ antibody POM2 as described above. Western blots were quantified on native photographs (uncropped, naïve images have been deposited on FigShare https://doi.org/10.6084/m9.figshare.11940606), representative western blot images in main and supplementary figures have been processed (linear adjustment of contrast and brightness) for better visualization.

### SPR

The binding properties of the complexes between rmPrP, POM1 and pomologs were measured at 298K on a ProteOn XPR-36 instrument (Bio-Rad) using 20mM HEPES pH 7.2 150mM NaCl 3mM EDTA and 0.005% Tween-20 as running buffer. mPrP was immobilized on the surface of GLC sensor chips through standard amide coupling. Serial dilution of antibodies (full IgG, Fab or single chain versions) in the nanomolar range were injected at a flow rate of 100 μL/min (contact time 6 minutes); dissociation was followed for 5 minutes. Analyte responses were corrected for unspecific binding and buffer responses by subtracting the signal of both a channel where no PrP was immobilized and a channel were no antibody was added. Curve fitting and data analysis were performed with Bio-Rad ProteOn Manager software (version 3.1.0.6).

### Statistical analyses

All data are given as mean, error bars represent standard deviation (Fig. S14C) or standard error of the mean (Fig. S20). The exact sample sizes and test details are given for each graph in the Supplementary Table 2. All biological measurements are taken from distinct samples. Statistical analysis and visualization were performed using Prism 8 (GraphPad).

### Synchrotron radiation circular dichroism (SRCD)

Secondary structure content of complexes between rmPrP and POM1, ^hc^Y57 and ^hc^Y104A was analyzed with Synchrotron radiation circular dichroism (SRCD) spectroscopy.

Experiments were performed using a nitrogen-flushed B23 beamline for synchrotron radiation circular dichroism (SRCD) at Diamond Light Source or ChirascanPlus CD spectropolarimeter (Applied Photophysics Ltd, Leatherhead, UK). With both instruments, scans were acquired at 20°C using an integration time of 1 sec and 1 nm bandwidth. Demountable cuvette cells of 0.00335 cm pathlength were used in the far-UV region (180-260 nm) to measure the CD of the protein concentration varying from 10 to 102 μM of proteins in 10 mM NaP pH 7; 150 mM NaCl. Mixtures were prepared to a stoichiometric molar ratio of 1:1. SRCD data were processed using CDApps (*32*) and OriginLab™. Spectra have been normalized using average amino acid molecular weight of 113 for secondary structure estimation from SRCD and CD spectra was carried out using CDApps using the Continll algorithm (*33*). For comparison of calculated and observed spectra, full molecular weight of sample and complex were used. Measurement on free mPrP and free antibodies was done as reference.

## Notes

https://doi.org/10.6084/m9.figshare.11940606

### Competing Interest Statement

The authors have declared no competing interest.

