## Supplementary Figure S1 for "A conformational switch controlling the toxicity of the prion protein"

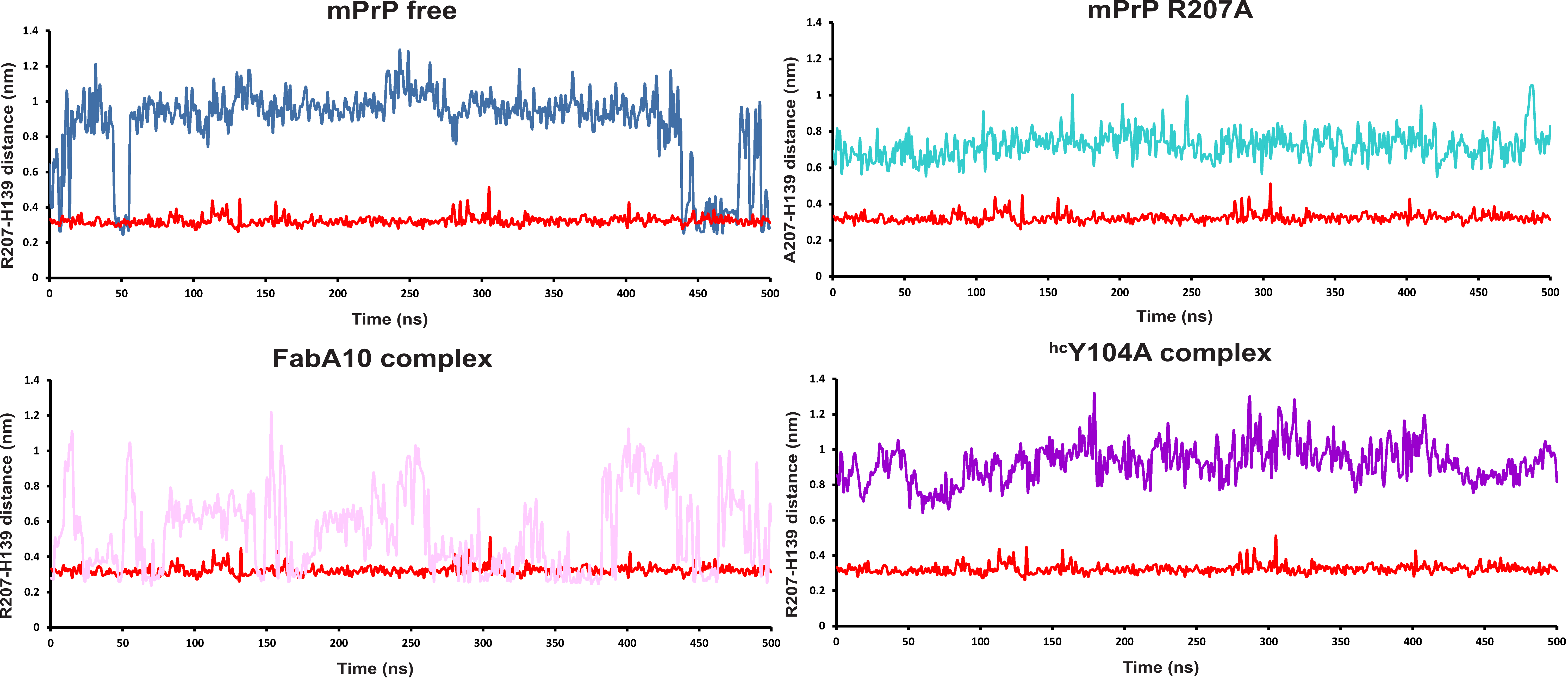


***Supplementary Figure 1***. Distances between the R207 and H139 residues in MD simulations of PrP-Ab complexes. The simulation of the PrP-POM1 complex (red) in reproduced in all charts to facilitate comparisons. When PrP is bound to POM1, the H-bond between R207 and H139 (termed H-latch) is always present, with distance between the centroid of their sidechains around 0.3 nm. Greater distances indicate loss of hydrogen interactions and consequently absence of the H-latch. The complex of PrP with the $^{hc}$Y104A pomolog never shows formation of the H-latch, whereas FabA10 shows intermediate values. Simulations were run in triplicate but only representative traces are shown; aggregated analyses are shown in Fig 1B.
