## Supplementary Figure S2 for "A conformational switch controlling the toxicity of the prion protein"

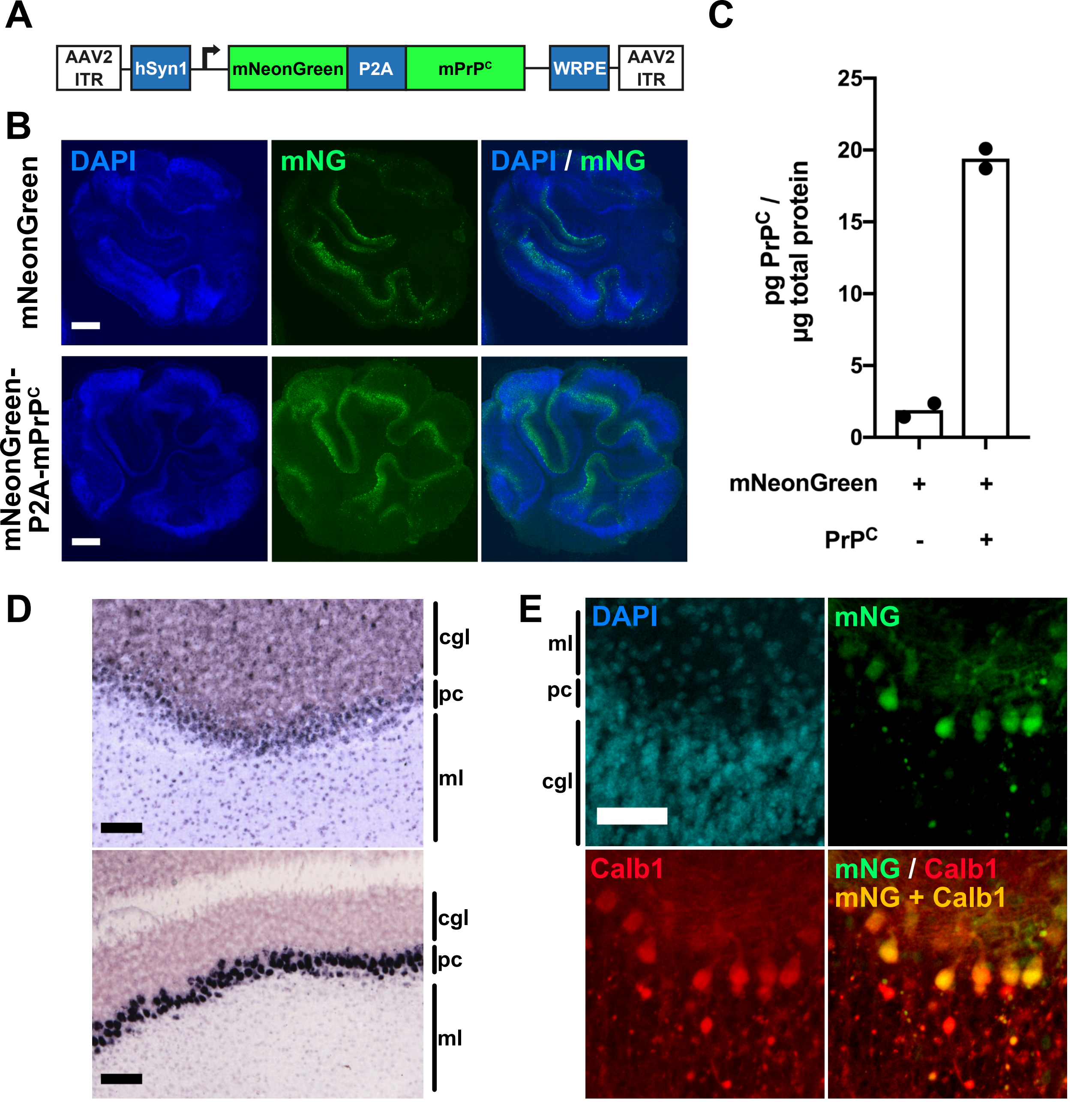


***Supplementary Figure* 2**. **(A)** Scheme of AAV used for bi-cistronic expression of monomeric NeonGreen and PrP$^{C}$, separated by a P2A site (mNG-P2A-PrP$^{C}$, scissors indicate ribosomal skipping site). hSyn1 = human Synapsin 1 promoter. WRPE = woodchuck hepatitis virus regulatory posttranscriptional element. ITR = inverted terminal repeats. **(B)** Robust expression of mNG-P2A-PrP$^{C}$ on fluorescent micrographs from transduced *Prnp*$^{ZH3/ZH3}$ COCS. Scale bar = 500 µm **(C)** Holo-POM19 / holo-POM2-biotin PrP$^{C}$ sandwich ELISA of samples depicted in (B). **(D)** Representative images of expression levels of Synapsin 1 (Syn1, *upper*) and Calbindin 1 (Calb1, *lower*) show predominant (Syn1) or almost exclusive (Calb1) expression in Purkinje cells (pc) in the cerebellar cortex. Image credit: Allen Institute. Scale bar = 100 µm. **(E)** Fluorescent micrographs of *Prnp*$^{ZH3/ZH3}$ COCS transduced with the AAV outlined in panel (A) show mNeonGreen expression predominantly in calbindin 1-expressing Purkinje cells. Scale bar = 50 µm. cgl = cerebellar internal granular layer, pc = Purkinje cell layer, ml = molecular layer.
