## Supplementary Figure S3 for "A conformational switch controlling the toxicity of the prion protein"

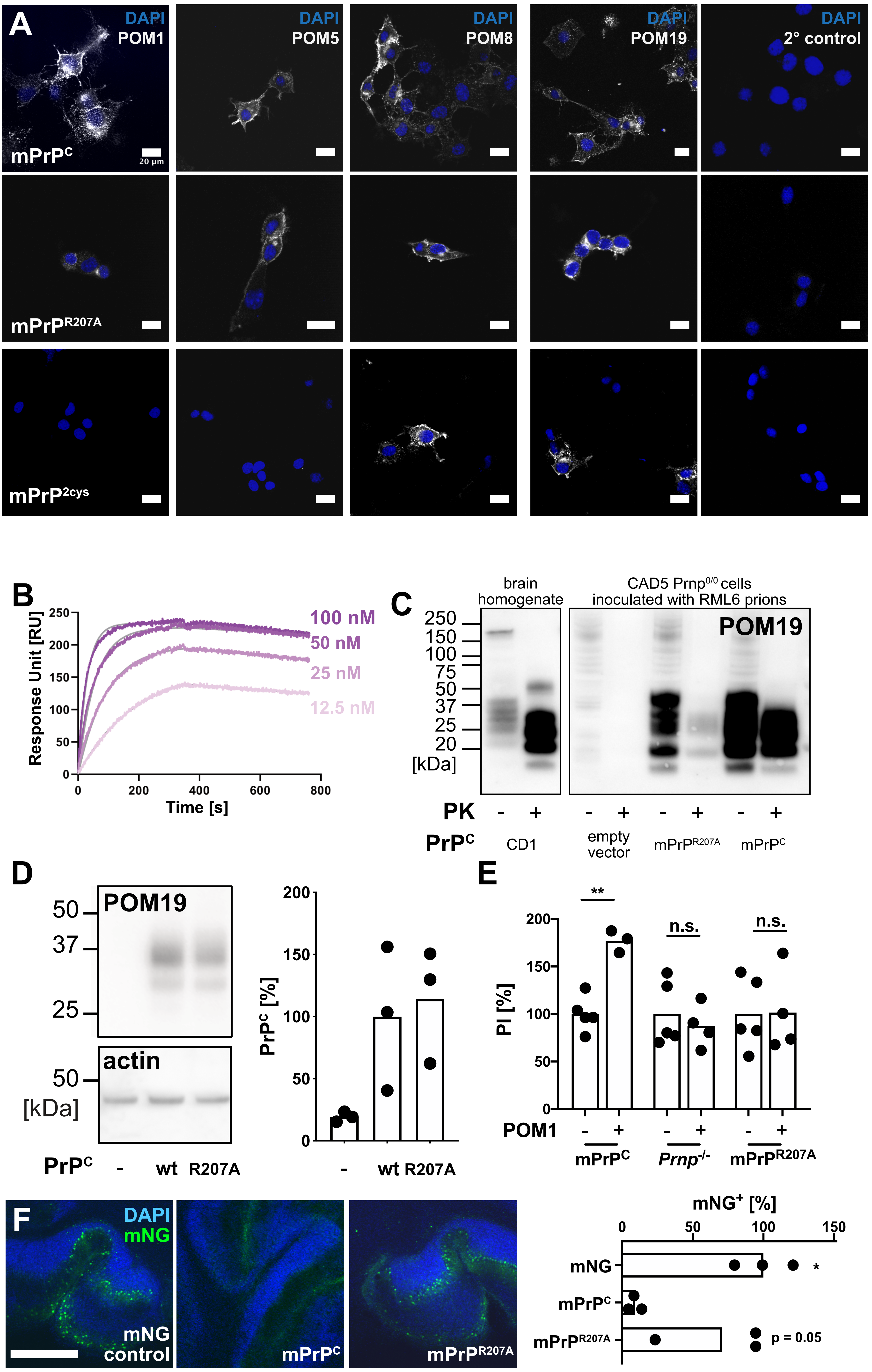


***Supplementary Figure 3.*** **(A)** Immunohistochemistry of CAD5 *Prnp*$^{-/-}$ cells stably transfected with pcDNA3.1 vector expressing wild-type murine PrP$^C$ (mPrP$^C$), mPrP$^{R207A}$ and mPrP$^{2cys}$. Monoclonal anti-PrP$^C$ antibodies targeting distinct conformational epitopes on the globular domain of PrP$^C$ were incubated to assess conformational changes in mPrP$^{R207A}$ (POM1: α1-α3, POM5: β2-α2, POM8: α1-α2, POM19: β1-α3). Except for diminished staining of POM1 in mPrP$^{R207A}$, we observed robust detection of mPrP$^{R207A}$ by POM5, POM8 and POM19 and mPrP$^{2cys}$ by POM8 and POM19. Scale bar = 20 µm. **(B)** Surface plasmon resonance (SPR) traces showing binding of POM1 to recombinant mPrP$^{R207A}$ (rmPrP$^{R207A}$, $k_a$=3.8E$^{+05}$ 1/Ms, $k_d$=1.8E$^{-04}$ 1/s, $K_D$=4.7E$^{-10}$ M; for comparison binding to recombinant wild-type murine PrP showed $k_a$=3.6E$^{+05}$ 1/Ms; $k_d$=9.1E$^{-05}$ 1/s ; $K_D$=2.5E$^{-10}$ M). **(C)** Proteinase K digestion of brain homogenates and cell lysates from chronically RML6-inoculated CAD5 cells (4$^{th}$ passage is shown) detected with POM19. RML6 prions (*lanes 1 and 2*) and inoculated CAD5-mPrP$^C$ cells (*lanes 7 and 8*) show a typical "diagnostic shift" of PK-digested PrP$^{Sc}$, whereas only trace amounts of PrP$^{Sc}$ are detectable in CAD5-mPrP$^{R207A}$ cells (*lanes 5 and 6*). Lack of detectable PrP$^{Sc}$ in CAD5 *Prnp*$^{-/-}$ (*lanes 3 and 4*) indicates no residual inoculum. Lanes are from non-adjacent samples blotted on the same membrane. Uncropped western blots can be found in auxillary supplementary materials **(D)** Stably transfected CAD5-mPrP$^C$ and CAD5-mPrP$^{R207A}$ cells show similar PrP$^C$ expression levels. *Right panel*: POM19 immunoreactivity is divided by actin immunoreactivity, values are given as percentages of PrP$^C$. **(E)** Addition of POM1 causes toxicity to CAD5 cells (left) but not to *Prnp*$^{-/-}$ or mPrP$^{R207A}$ CAD5 (*center and right*). The percentage of propidium iodide positive cells, determined by FACS, is shown on the y-axis. Values are given as percentages of CAD5 mPrP$^C$ PI-positive cells without POM1. n.s. not significant, ** $p < 0.01$ **(F)** *Prnp*$^{ZH3/ZH3}$ COCS transduced with wild-type mPrP$^C$ are susceptible to POM1 toxicity whereas COCS transduced with control vector ("mNG control") or mPrP$^{R207A}$ are not. Inserts highlight cell loss in mPrP$^C$ COCS. Values are given as percentage of empty control. * $p < 0.05$, ** $p < 0.01$ (mPrP$^C$ versus other groups). Scale bar: 500 µm
