## Supplementary Figure S4 for "A conformational switch controlling the toxicity of the prion protein"

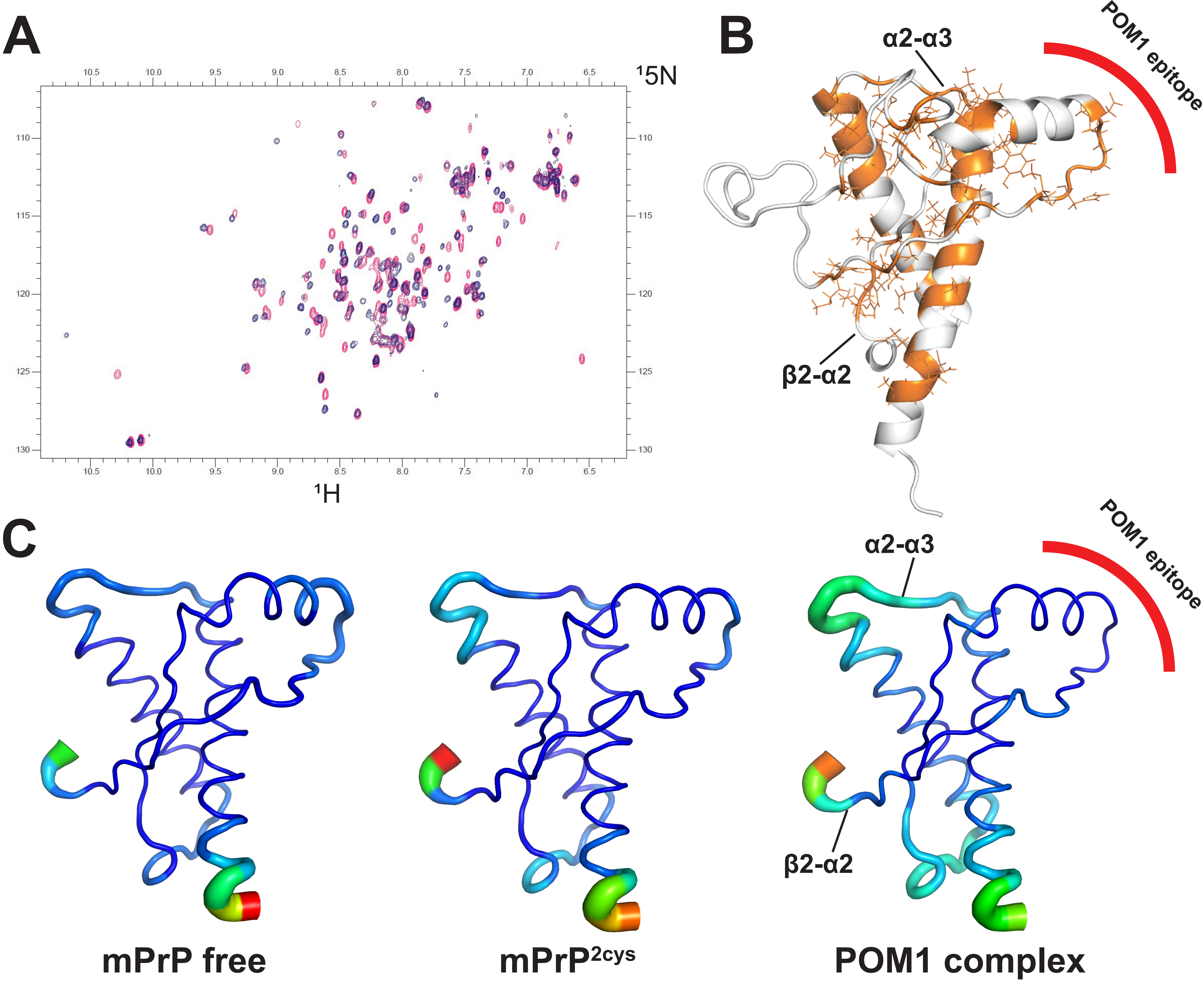


***Supplementary figure 4.*** (**A**) $^{15}$N-HSQC spectra of rmPrP free (red) and mPrP$^{2cys}$ (blue). Residues with different chemical shift in the two spectra are colored orange on the GD structure in **(B)**, which resemble the H-latch conformation in the POM1-PrP complex.(**C)** Molecular Dynamics simulations show that mPrP$^{2cys}$ resembles the PrP-POM1 complex, with increased flexibility in α2-α3 and β2-α2 loop and decreased flexibility in the 2Cys region, corresponding to the POM1 epitope.
