## Supplementary Figure S5 for "A conformational switch controlling the toxicity of the prion protein"

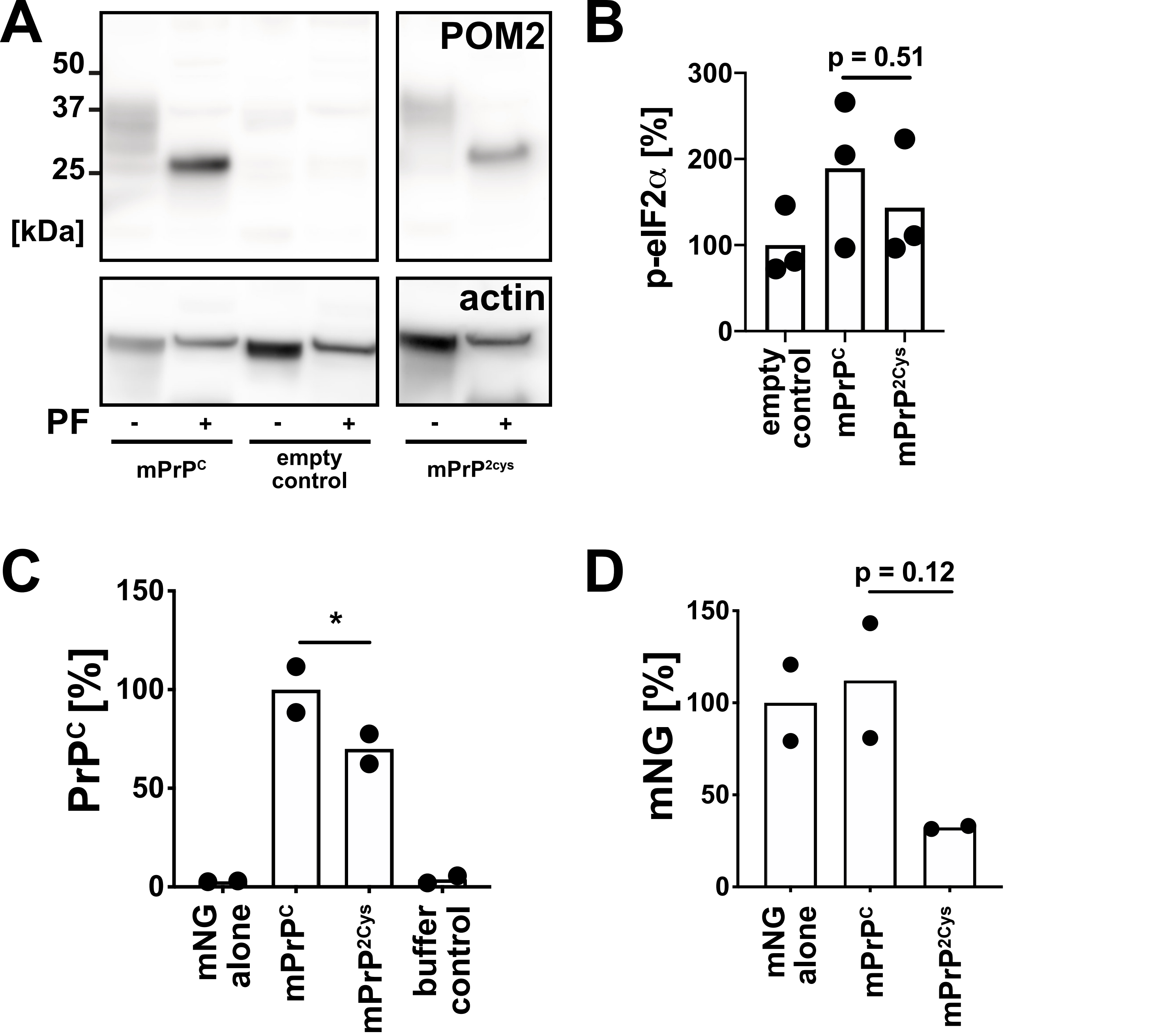


***Supplementary Figure* 5. (A)** PNGase-F digestion of cell lysates induced a shift in both murine wild-type $PrP^C$ and $mPrP^{2cys}$, indicating that both moieties had undergone N-linked glycosylation to a similar extent. Non-adjacent lanes were merged from the same gel. **(B)** CAD5 *Prnp*$^{-/-}$ cells expressing $mPrP^{2cys}$ did not show an upregulation of the unfolded protein response, suggesting that $mPrP^{2cys}$ did not undergo pathological degradation. Values are given as percentage of empty control vector (p-eIF2α / eIF2α / actin). **(C)** A POM2/POM3 sandwich ELISA of COCS transduced with empty control, $mPrP^C$ , $mPrP^{2cys}$ and buffer control shows robust $mPrP^{2cys}$ expression in transduced COCS, albeit significantly less than wild-type $mPrP^C$. Slices were harvested at 28 days post-transduction. **(D)** Reduced levels of mNG in *Prnp*$^{-/-}$ (ZH3) COCS expressing $mPrP^{2cys}$. mNG immunoreactivity values are divided by actin immuno-reactivity and expressed as percentages of empty control. Slices were harvested at 28 days post-transduction. * $p < 0.05$. Raw, uncropped blots can be found in Extended Supplementary Methods.
