## Supplementary Figure S6 for "A conformational switch controlling the toxicity of the prion protein"

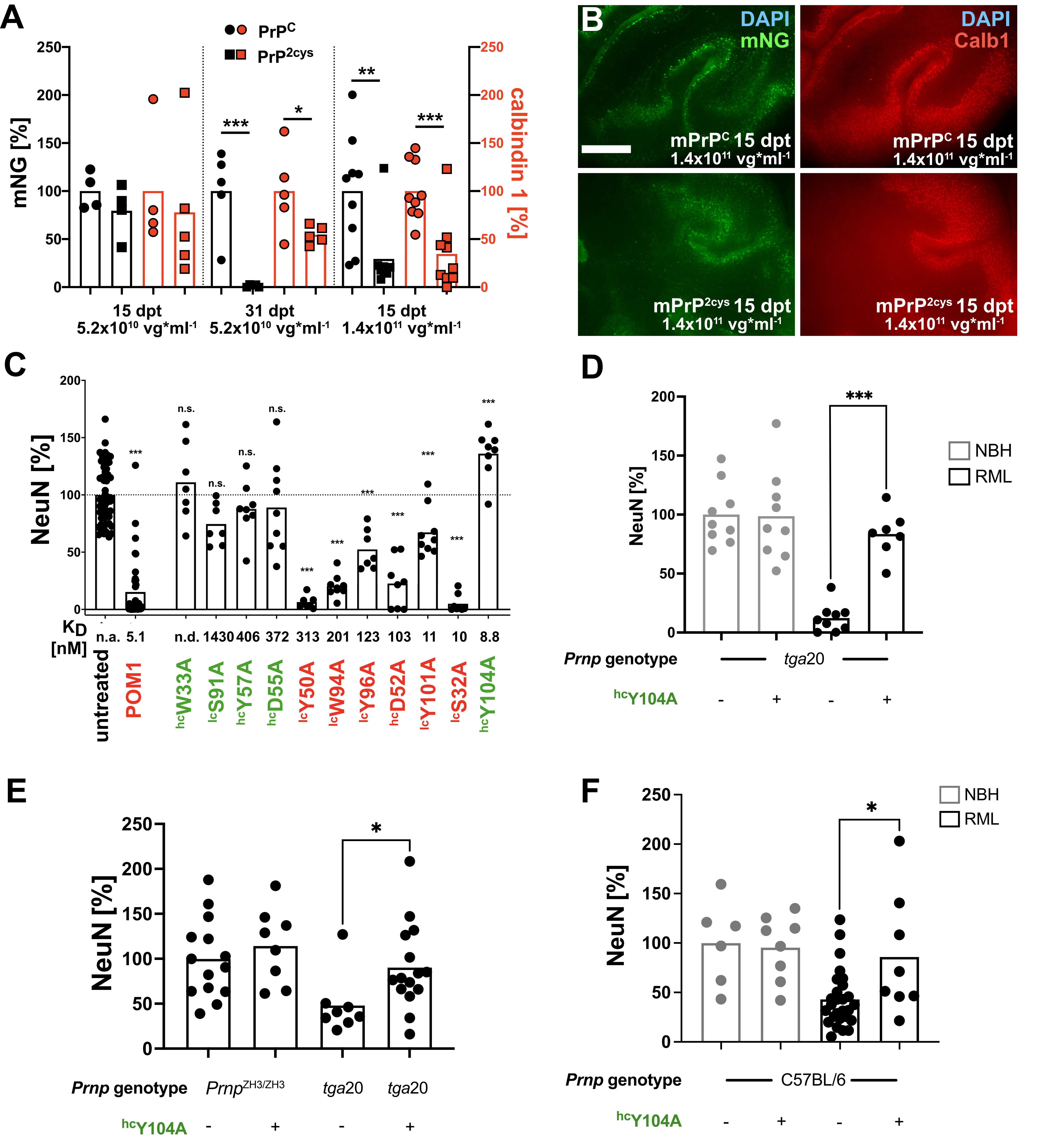


***Supplementary Figure* 6.** **(A)** Quantification of mNG and Calb1 fluorescence intensity from experiments shown in Fig. 2A+B and panel **(B)**. Values are given as percentages of mPrP$^{C}$. **(B)** Dose escalation of 2 x more viral genomes compared to Fig. 2A+B, lead to earlier onset of mPrP$^{2cys}$-mediated neurodegeneration. Significant neurodegeneration was observable as 15 days post-transduction. Scale bar: 500 µm. **(C)** Morphometric quantification of *tga*20 COCS treated with pomologs, representative photomicrographs are depicted in Fig. 2C. Graph shows aggregated data from multiple experiments performed under identical conditions. Innocuous pomologs are highlighted in green, POM1 and toxic pomologs are highlighted in red. 100% = untreated COCS, comparison of untreated COCS against all other groups. **(D)** Morphometric quantification of experiments depicted in Fig. 2D. 100% untreated COCS inoculated with NBH. Untreated, RML-infected COCS were compared against treated, RML-infected COCS. **(E)** Morphometric quantification of experiments depicted in Fig. 2E. 100% = untreated *Prnp*$^{ZH3/ZH3}$ COCS inoculated with 22L. NeuN fluorescence of untreated, 22L-infected *tga*20 COCS was compared against all other COCS. **(F)** Morphometric quantification of experiments depicted in Fig. 2F. 100% = untreated COCS inoculated with NBH. NeuN fluorescence of untreated, RML-infected COCS was compared against all other COCS. n.s. not significant, *** $p < 0.001$, ** $p < 0.01$, * $p < 0.05$.
