## Supplementary Figure S7 for "A conformational switch controlling the toxicity of the prion protein"

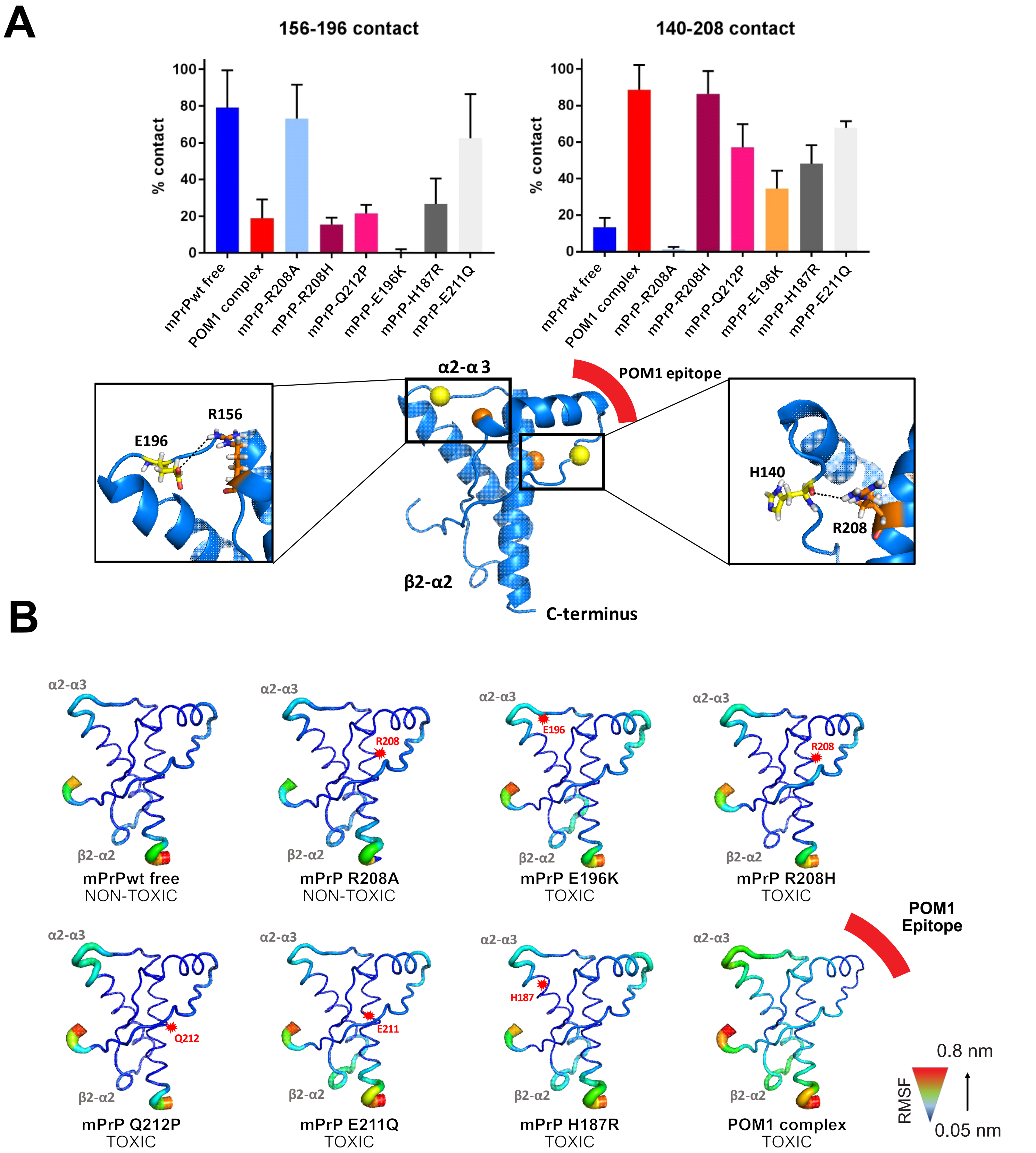


**(A)** MD simulations of POM1 binding and pathogenic *PRNP* mutations causing genetic prion disease show the R156-E196 interaction is abolished and induction of the H140-R208 H-latch is established,
**(B)** In agreement with this view, POM1 and human, hereditary PrP mutations responsible for fatal prion diseases favor altered flexibility in the α2-α3 and β2-α2 loop.
