## Supplementary Figure S8 for "A conformational switch controlling the toxicity of the prion protein"

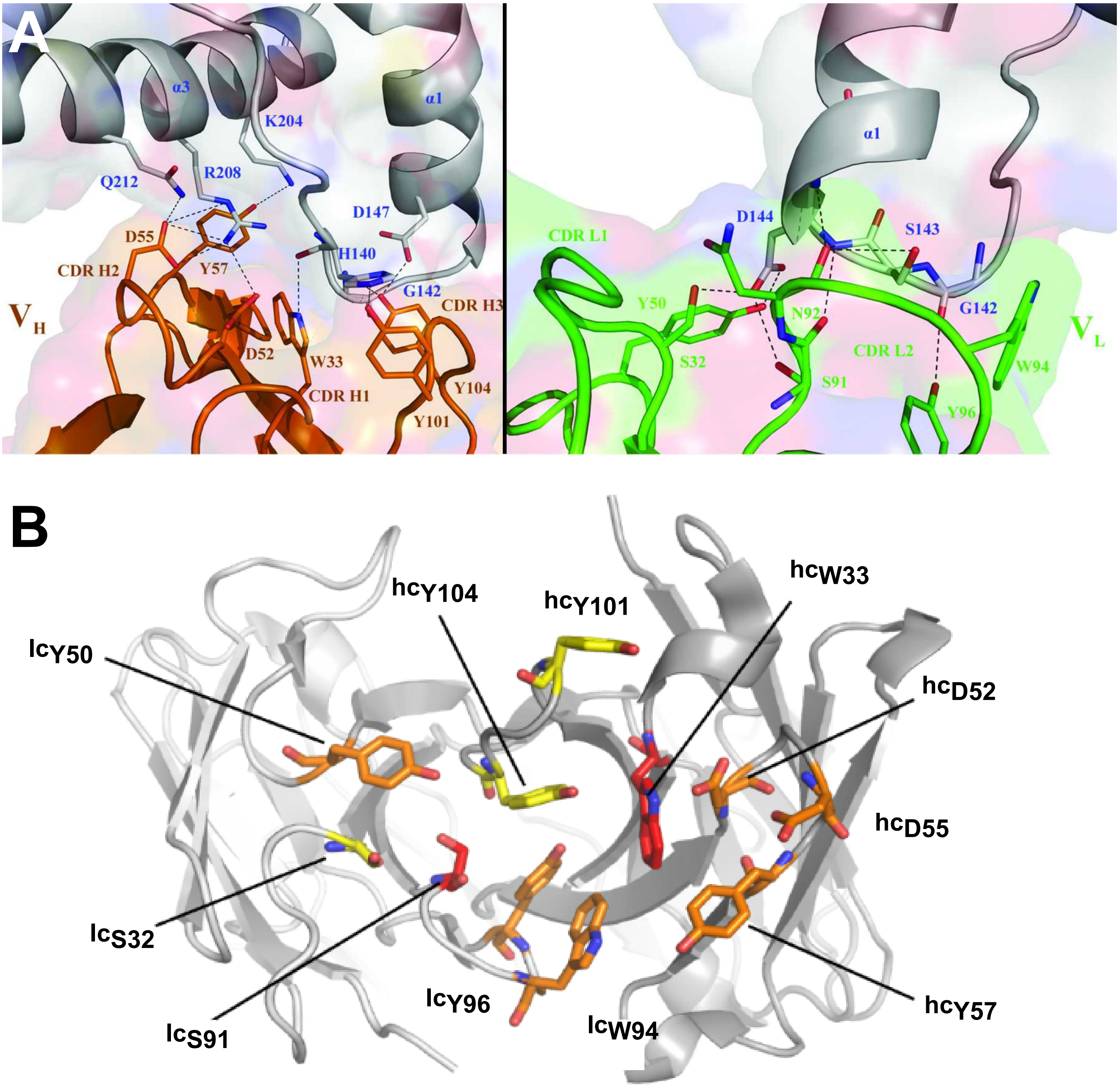


***Supplementary Figure 8.*** **(A)** Intermolecular contacts between human $PrP^{C}_{120\text{-}230}$ and POM1 Fab variable heavy chain (magenta, *left panel*), and POM1 Fab variable light chain (green, r*ight panel*) as determined by Baral et al., 2012 (*8*). Reproduced with permission of the International Union of Crystallography from doi:10.1107/S0907444912037328. **(B)** Schematic representation of a single-chain fragment of wild-type POM. The mutated residues are indicated as stick on the cartoon structure of POM1, color coded as in Supplementary Table 1B. The CDR loops are shown from the perspective of the antigen.
