## Supplementary Figure S9 for "A conformational switch controlling the toxicity of the prion protein"

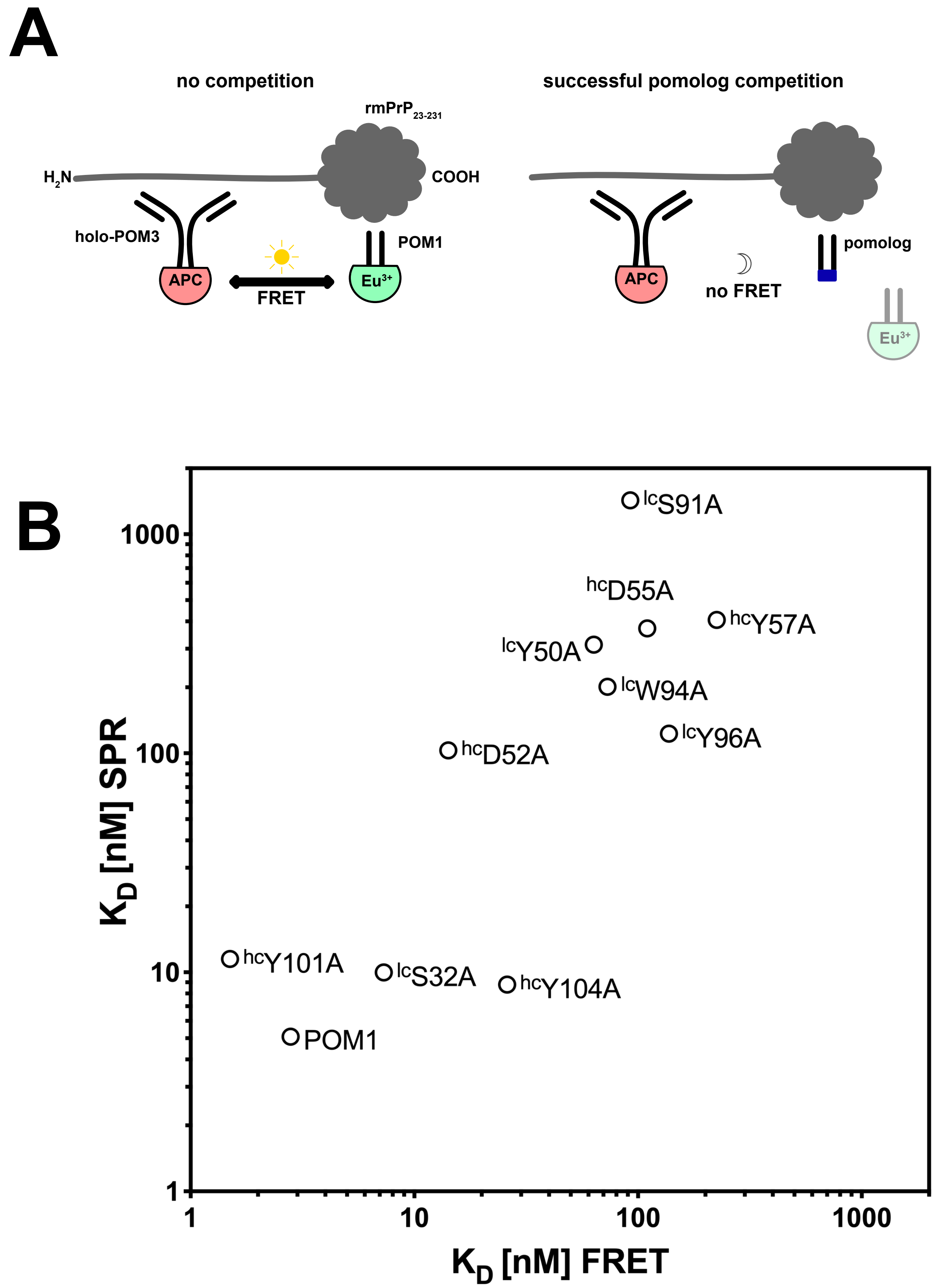


***Supplementary Figure* 9. (A)** Scheme of competition FRET assay to assess the $K_D$ of various pomologs. In the absence of competing antibody, FRET occurs due to proximity of allophycocyanin (APC)-labeled holo-POM3 and europium (Eu3+)-labeled POM1 (*left panel*). Because of liquid-phase competition, addition of unlabeled pomologs leads to a decrease in FRET signal (*right panel*). The calculation of binding constants from FRET is detailed in *Supplementary Methods*. **(B)** The binding constants measured by SPR and by FRET were in good agreement, Spearman r = 0.77, p<0.01, 95% CI 0.30-0.94) with the exception of $^{hc}$W33A, whose binding on SPR was too weak to be precisely measured.
