## Supplementary Figure S10 for "A conformational switch controlling the toxicity of the prion protein"

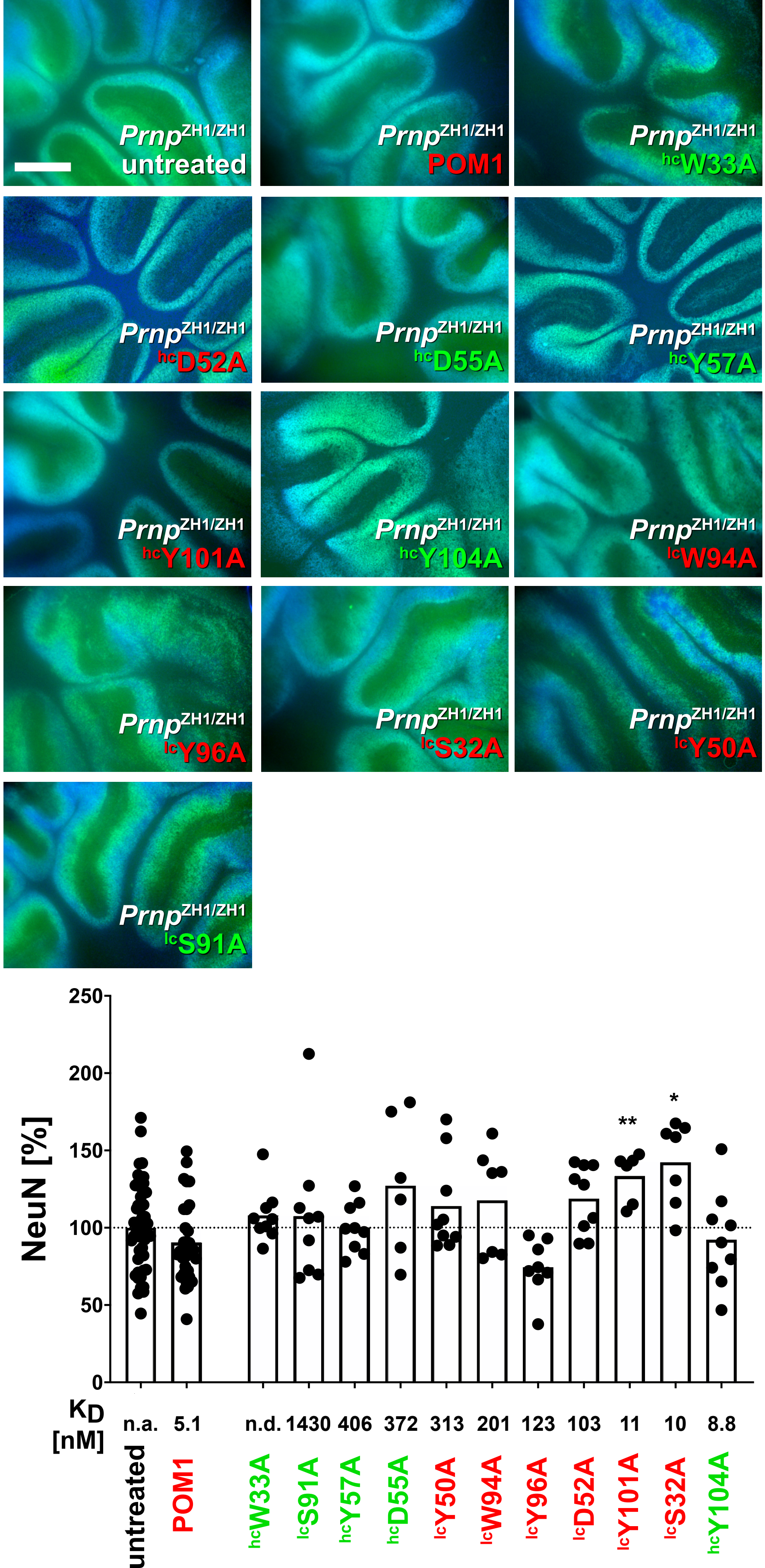


***Supplementary Figure* 10.** Representative fluorescent micrographs of *Prnp*$^{ZH1/ZH1}$ COCS treated with pomologs and morphometric quantification. In all cases, neuronal viability is similar to that of controls, indicating that neither POM1 nor any of the pomologs exert a toxic effect onto PrP-deficient tissue. Innocuous / protective pomologs are colored green, toxic pomologs red. Graph shows aggregated data from multiple experiments performed under identical conditions. NeuN fluorescence from untreated *Prnp*$^{ZH1/ZH1}$ COCS = 100%. Scale bar: 500 µm. ** $p < 0.01$, * $p < 0.05$
