## Supplementary Figure S11 for "A conformational switch controlling the toxicity of the prion protein"

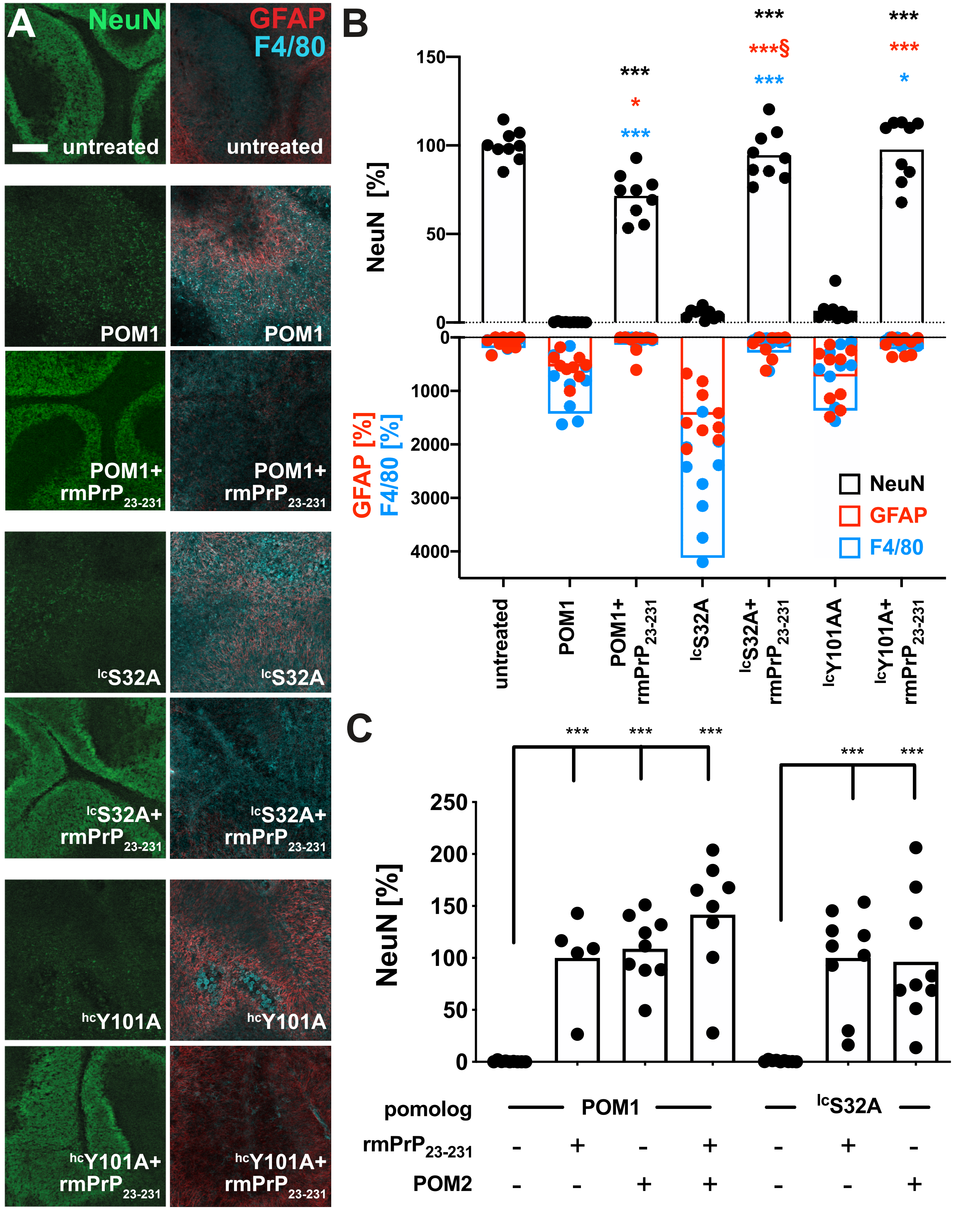


***Supplementary Figure* 11. (A)** Immunofluorescent stainings of COCS with NeuN (neurons), GFAP (astrocytes) and F-4/80 (microglia and macrophages). The high-affinity toxic pomologs lcS32A and hcY101A were administered to *tga20* COCS, either alone or after preincubation with equimolar amounts of soluble $rmPrP_{23\text{-}231}$. POM1, lcS32A and lcY101A induced profound neuronal loss, astrogliosis and activation of microglia, which was completely prevented by pre-incubation with their cognate antigen. **(B)** Morphometric quantification of fluoresence intensity from images depicted in panel A. §: one extreme outlier was excluded from the analysis (y=2046.3%, $p<0.05$, extreme studentized deviate method). Values are given as percentages of untreated control. Pairwise comparison of pomolog in the presence or absence of $rmPrP_{23\text{-}231}$. **(C)** Pre-incubation of the toxic, high-affinity pomolog lcS32A with the $PrP^C$ octarepeat binder POM2 led to ablation of neurotoxicity. Values are normalized to POM1+$rmPrP_{23\text{-}231}$ (*bars 1-4*) or lcS32+$rmPrP_{23\text{-}231}$ (*bars 5-7*). *** $p<0.001$, ** $p<0.01$, * $p<0.05$.
