## Supplementary Figure S12 for "A conformational switch controlling the toxicity of the prion protein"

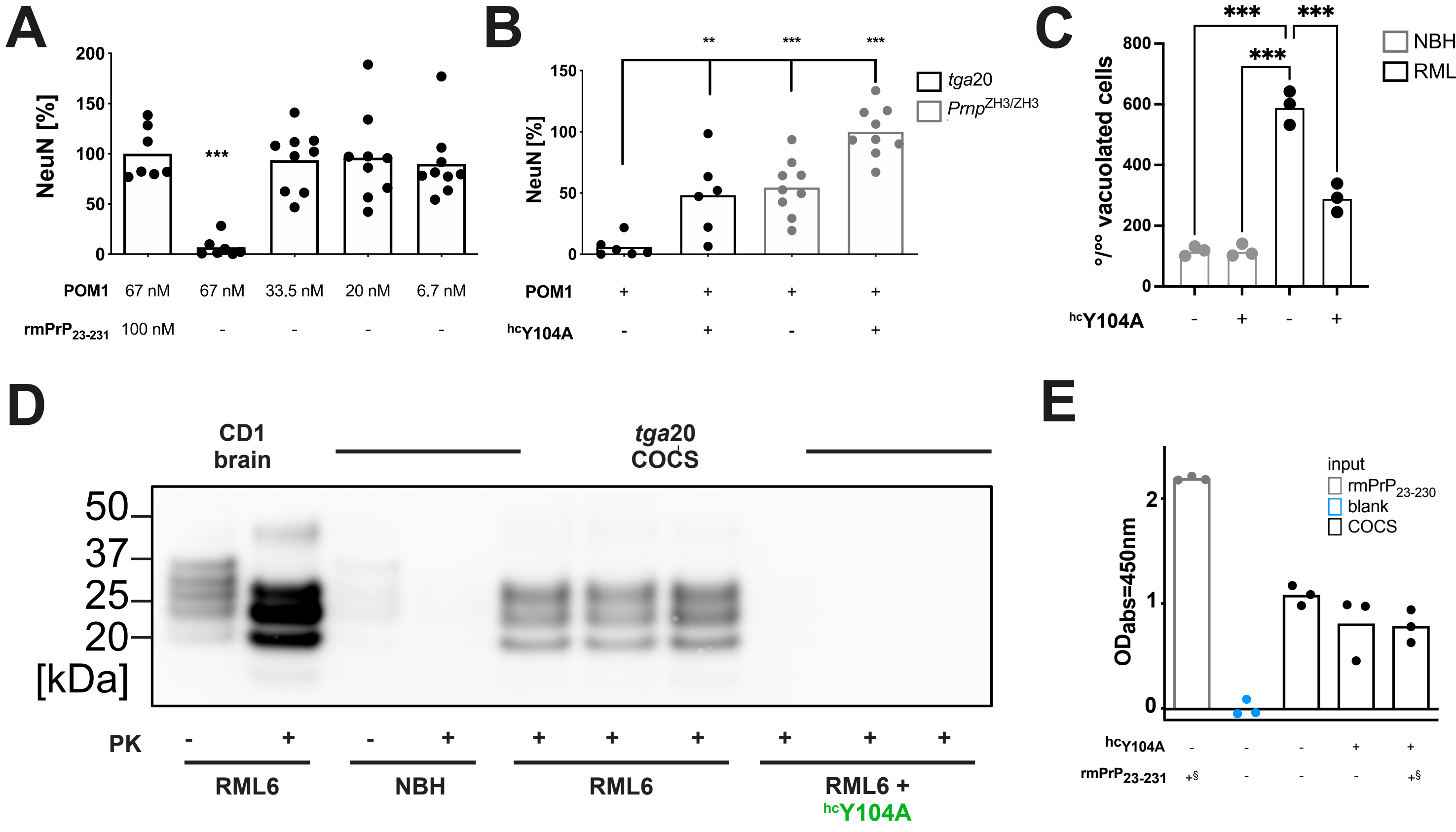


***Supplementary Figure*** **12. (A)** The minimal toxic dosage of POM1 was determined to be 67 nM for 14 days in *tga*20 COCS. Values are given as percentages of POM1 + $rmPrP_{23\text{-}231}$. **(B)** Pre-incubation of *tga*20 COCS with 1.2 µM of $^{hc}$Y104A prevented POM1-induced toxicity. Values are given as percentages of $Prnp^{0/0}$ COCS treated with POM1 + $^{hc}$Y104A. **(C)** Treatment of *tga*20 COCS with $^{hc}$Y104A for 7 days did not reduce $PrP^{C}$ levels, as determined by $PrP^{C}$ sandwich ELISA. § 870 pM of $rmPrP_{23230}$ were used as positive control (*first lane*). Pomologs were pre-incubated with 600 nM of $rmPrP_{23\text{-}230}$ as negative controls (*last lane*). Ordinate: absorbance, given as optical density at λ = 450 nm **(D** Treatment with $^{hc}$Y104A (180nM; 5 days) reduced vacuolation in chronically prion-infected Gt1 cells. Each dot represents a separate experiment (1000 cells/experiment, chi-square test). **(E)** Treatment of prion-infected *tga*20 COCS with $^{hc}$Y104A led to a reduction in $PrP^{Sc}$ levels. 1 lane corresponds to a pool of 6-9 COCS digested with proteinase K (PK), $PrP^{Sc}$ was detected using holo-POM1. All graphs: *** $p < 0.001$, ** $p < 0.01$, * $p < 0.05$
