## Supplementary Figure S13 for "A conformational switch controlling the toxicity of the prion protein"

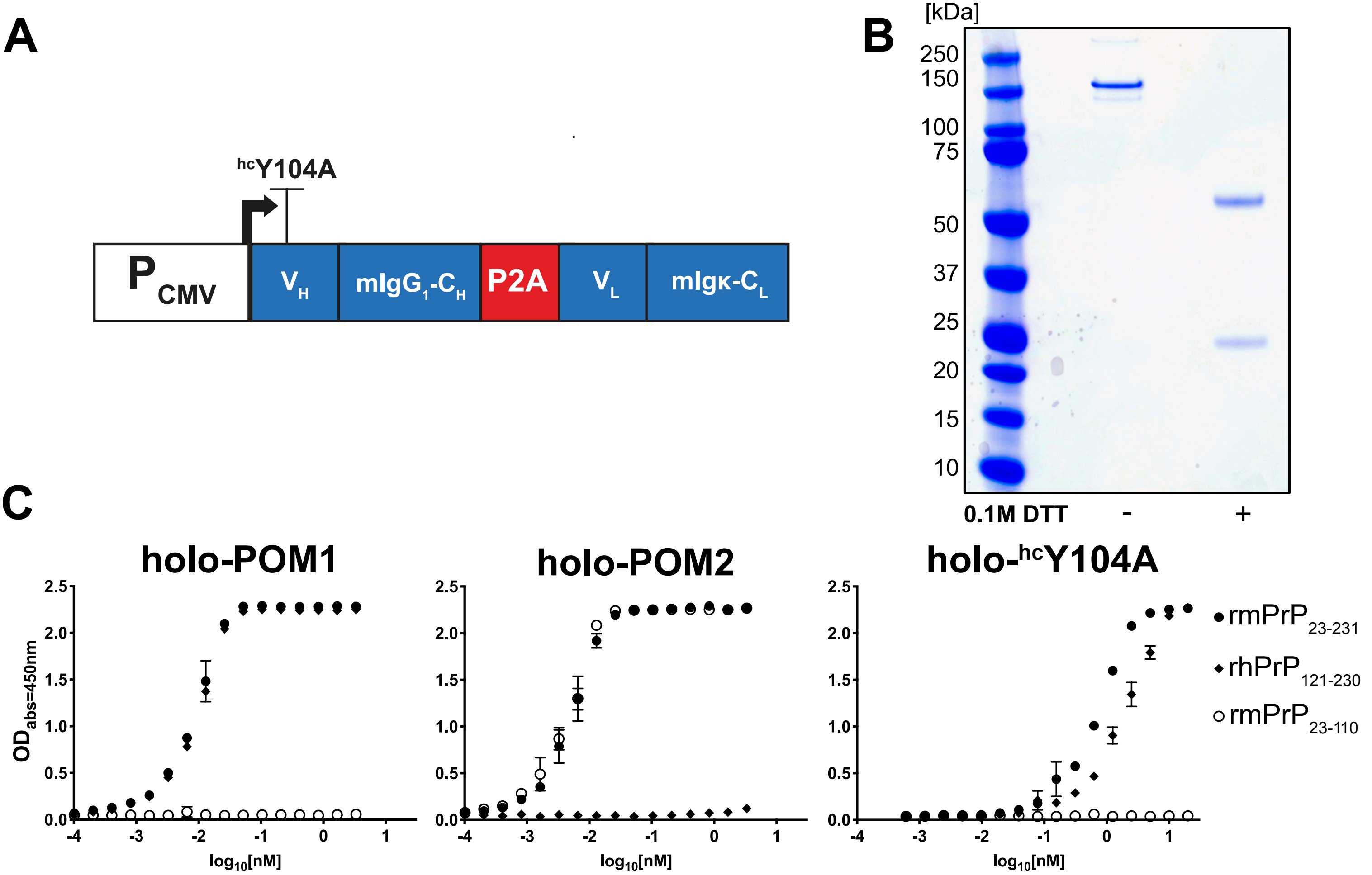


***Supplementary Figure*** **13**. **(A)** Schematic depiction of the vector for recombinant mammalian expression of holo-hcY104A. A bicistronic expression cassette encoding the variable domains of hcY104A were grafted onto a murine IgG1 and Igκ backbone driven by a cytomegalovirus promoter. A P2A self-cleaving site at the 3' end of the heavy-chain fragment results in co-translational cleavage of the polyprotein. **(B)** SDS-PAGE of purified holo-hcY104A in the presence and absence of 0.1M Dithiothreitol (DTT). **(C)** Indirect ELISA of holo-hcY104A and holo-POM1 showing (sub-) nanomolar affinity to full-length, recombinant mouse PrP ($mPrP_{23\text{-}230}$) and a globular domain fragment of recombinant human PrP ($mPrP_{121\text{-}231}$). As a control, holo-wtPOM2, which targets the flexible tail, bound to both $rmPrP_{23\text{-}231}$ and $rmPrP_{23\text{-}110}$, but not to $rhPrP_{121\text{-}230}$. Neither holo-hcY104A nor holo-POM1 bind to $rmPrP_{23\text{-}110}$. Nanomolar antibody concentrations are plotted in $\log_{10}$ scales on the abscissa. Error bars indicate standard deviation.
