## Supplementary Figure S14 for "A conformational switch controlling the toxicity of the prion protein"

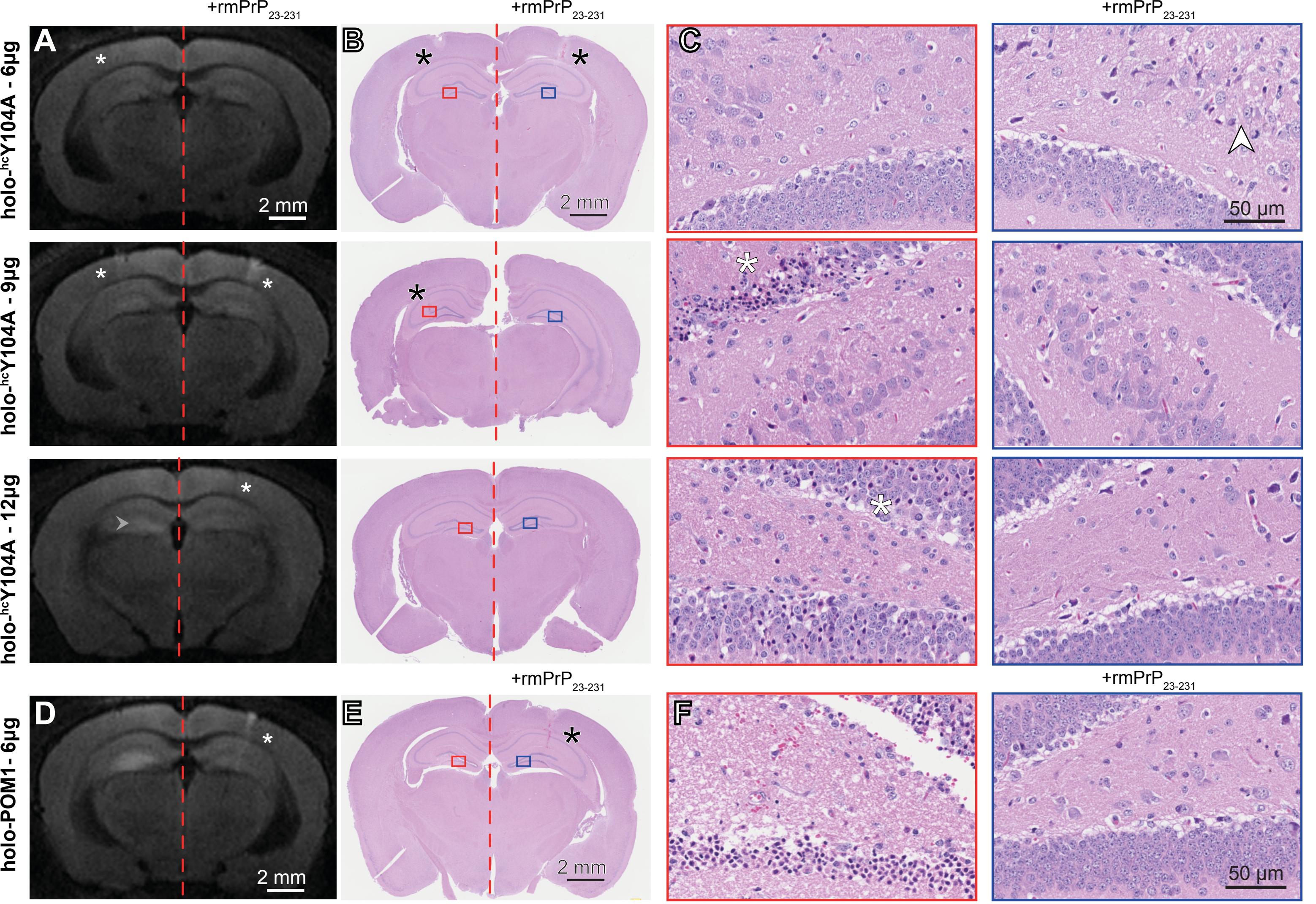


***Supplementary Figure* 14.** Representative magnetic resonance DWI images 24h after stereotactic injection of holo-$^{hc}$Y104A (*left*). Contralateral injections of holo-$^{hc}$Y104A + rmPrP$_{23231}$ (*right*). A small area of hyperintensity was found in one mouse after injection of 12 µg of holo-$^{hc}$Y104A (white arrowhead). *White asterisks*: needle tract. **(B)** Hematoxylin and eosin (HE) stained sections from mice shown in panel A. *Asterisks*: needle tract. Rectangles denote regions magnified in panel C. **(C)** HE sections (CA4). *Left*: holo-$^{hc}$Y104A injections (6, 9 and 12 µg). *Right*: holo-$^{hc}$Y104A+rmPrP$_{23\text{-}230}$. *Asterisk (9 µg)*: neurons with hypereosinophilic cytoplasm and nuclear condensation in the vicinity of the needle tract. *Asterisk (12 µg)*: These neurons were diffusely distributed among numerous healthy neurons. *White arrowhead*: vacuoles indicative of edema along the needle tract. **(D)** DWI images of 6 µg holo-POM1 +/- rmPrP$_{23\text{-}231}$, revealing a hyperintense signal at 24 hours. **(E)** Hematoxylin/eosin-stained section from a mouse shown in panel D. *Asterisks*: needle tract. *Rectangles*: areas in panel F. **(F)** HE sections (CA4). Holo-POM1 injections revealed damaged neurons with condensed chromatin and hypereosinophilc cytoplasm.
