## Supplementary Figure S15 for "A conformational switch controlling the toxicity of the prion protein"

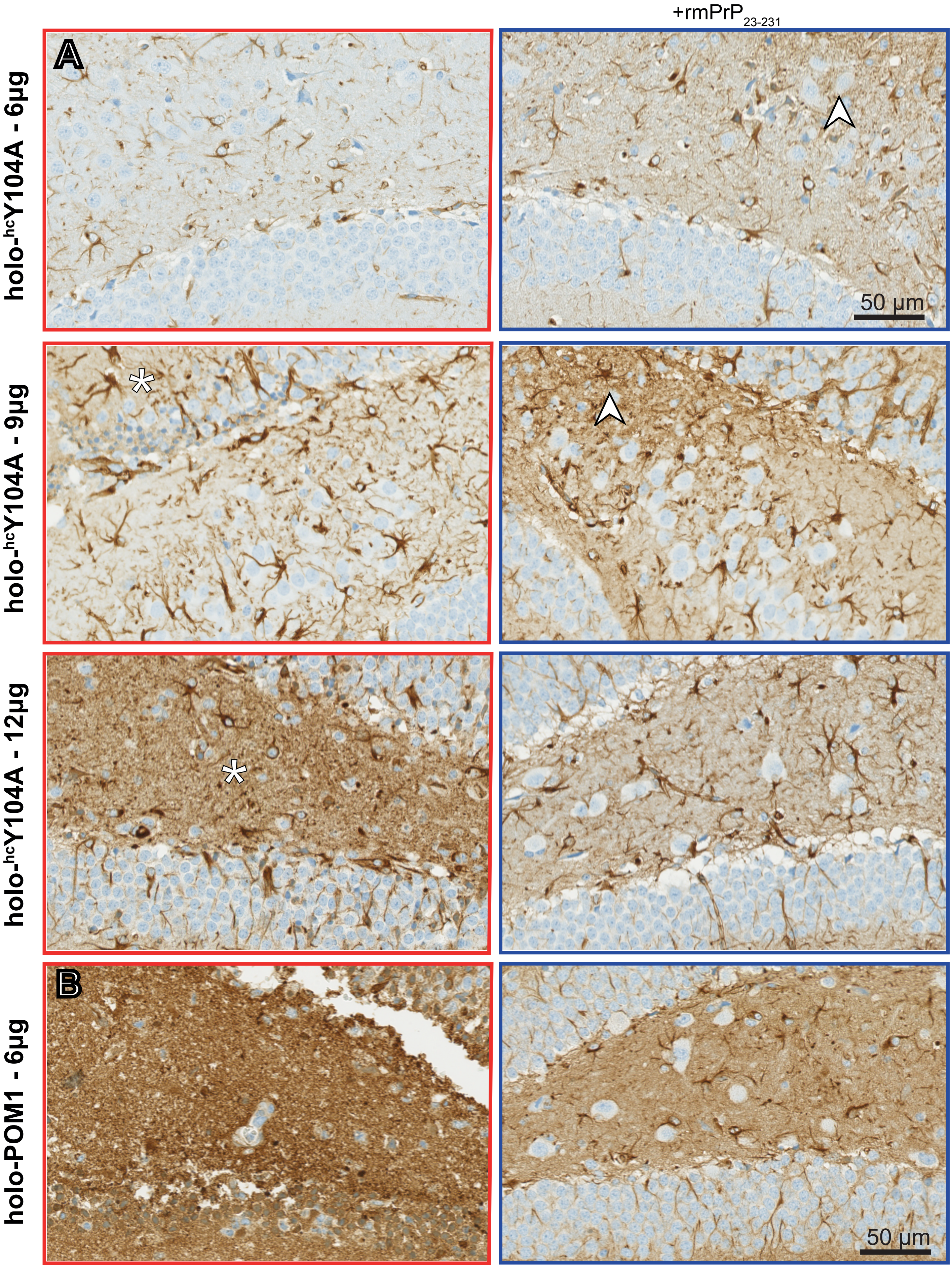


***Supplementary Figure 15***. (**A**) Photomicrographs of glial fibrillary acid protein (GFAP) immunohistochemistry on consecutive sections depicted in figure S15C. *Left column*: holo-$^{hc}$Y104A injections (6, 9 and 12 µg). *Right column*: holo-$^{hc}$Y104A + rmPrP$_{23\text{-}231}$. GFAP immunoreaction was increased in areas of neuronal damage (*white asterisks*) and around needle tracts (*white arrowheads*). (**B**) Micrographs demonstrating an intensive GFAP immunoreaction in areas with extensive holo-POM1 (6 µg)- induced neurotoxicity. *Left panel*: POM1 injection (6 µg). *Right panel*: holo-$^{hc}$Y104A + rmPrP$_{23\text{-}231}$. Sections are consecutive to those shown in figure S14F.
