## Supplementary Figure S16 for "A conformational switch controlling the toxicity of the prion protein"

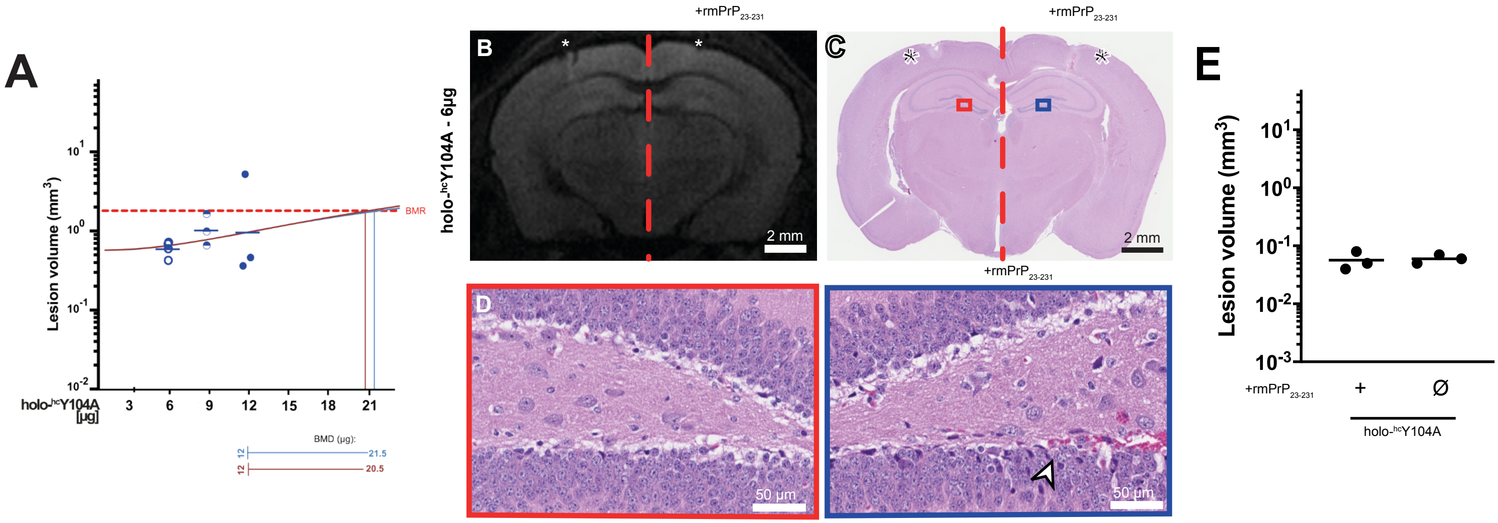


***Supplementary Figure* 16. (A)** A hypothetical benchmark dose analysis was performed using $log_{10}$-transformed lesion volumes corresponding to different amounts of holo-$^{hc}$Y104A (data from Fig. S14). BMR: Benchmark response (0.15 $mm^3$, dashed red line). The benchmark dose (BMD) is defined as the dose at the BMR. The vertical lines indicate the BMD values corresponding to the different dose response values (blue: 21.5 µg, brown line: 20.5 µg). The upper limit of the safe dose is provided by the lower 95% confidence interval of the BMD (horizontal lines below the graph: blue: 12 µg, brown: 12 µg). **(B)** Representative DWI images taken 24 h after stereotactic injection of 6 µg holo-$^{hc}$Y104A into male *tga*20 mice (*left half of the image*, injected into CA3). Contralateral side: 6 µg holo-$^{hc}$Y104A pre-incubated with an equimolar amount of $rmPrP_{23\text{-}230}$. *White asterisks*: needle tract. **(C)** Photomicrograph of HE-stained sections from mouse brain shown in panel B. *Asterisks*: needle tract. Rectangles correspond to regions magnified in panel C. **(D)** Higher magnification of the end-plate of the hippocampus. *Left panel*: holo-$^{hc}$Y104A. *Right panel*: holo-$^{hc}$Y104A preincubated with $rmPrP_{23\text{-}230}$. *Arrow*: needle tract. **(E)** Quantification of lesion volumes after injection of holo-$^{hc}$Y104A in contrast to control injection into *tga*20 mice (N = 3).
