## Supplementary Figure S17 for "A conformational switch controlling the toxicity of the prion protein"

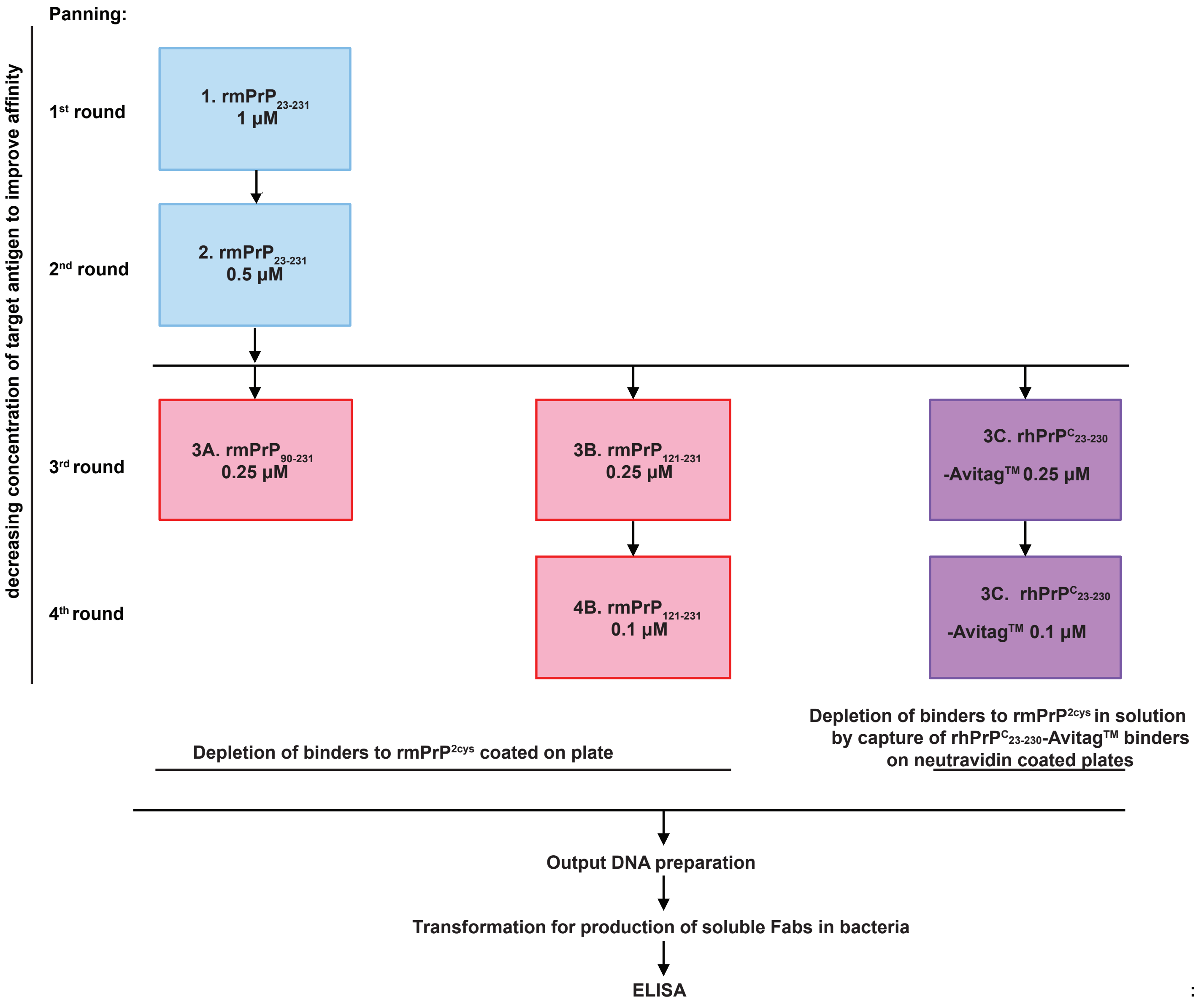


***Supplementary Figure* 17.** A synthetic human Fab phage library was used for panning. For each panning round, the targeted antigens are reported with the respective concentration. Full-length recombinant murine $PrP_{23\text{-}231}$ ($rmPrP_{23\text{-}231}$; light blue boxes) was used as a target for the first and the second round of phage panning. At the third and fourth round, phages were depleted of the binders to $rmPrP^{2cys}$ and selected for binding to either $rmPrP_{90\text{-}231}$ or $rmPrP_{121\text{-}231}$ (recombinant murine PrP fragments lacking the N-terminal flexible tail; light red boxes) or to recombinant human $PrP_{23\text{-}230}$-AviTag$^{TM}$ ($rhPrP_{23\text{-}230}$-AviTag$^{TM}$, purple boxes). In $rmPrP_{90\text{-}231}$ or $rmPrP_{121\text{-}231}$ panning, Fab-displayed Fab were depleted of binders to $rmPrP^{2cys}$ coated on plates. In $rhPrP_{23\text{-}230}$-AviTag$^{TM}$ panning, depletion of binders to $rmPrP^{2cys}$ in solution was achieved by capturing Fabs binding to $rhPrP_{23\text{-}230}$-AviTag$^{TM}$ on neutravidin coated wells. Polyclonal DNA preparation from the selected phages at the third round ($rmPrP_{90\text{-}231}$) and fourth round ($rmPrP_{121\text{-}231}$ and $rhPrP_{23\text{-}230}$-AviTag$^{TM}$) was used for transformation in bacteria and the screening of single clones by ELISA.
