## Supplementary Figure S18 for "A conformational switch controlling the toxicity of the prion protein"

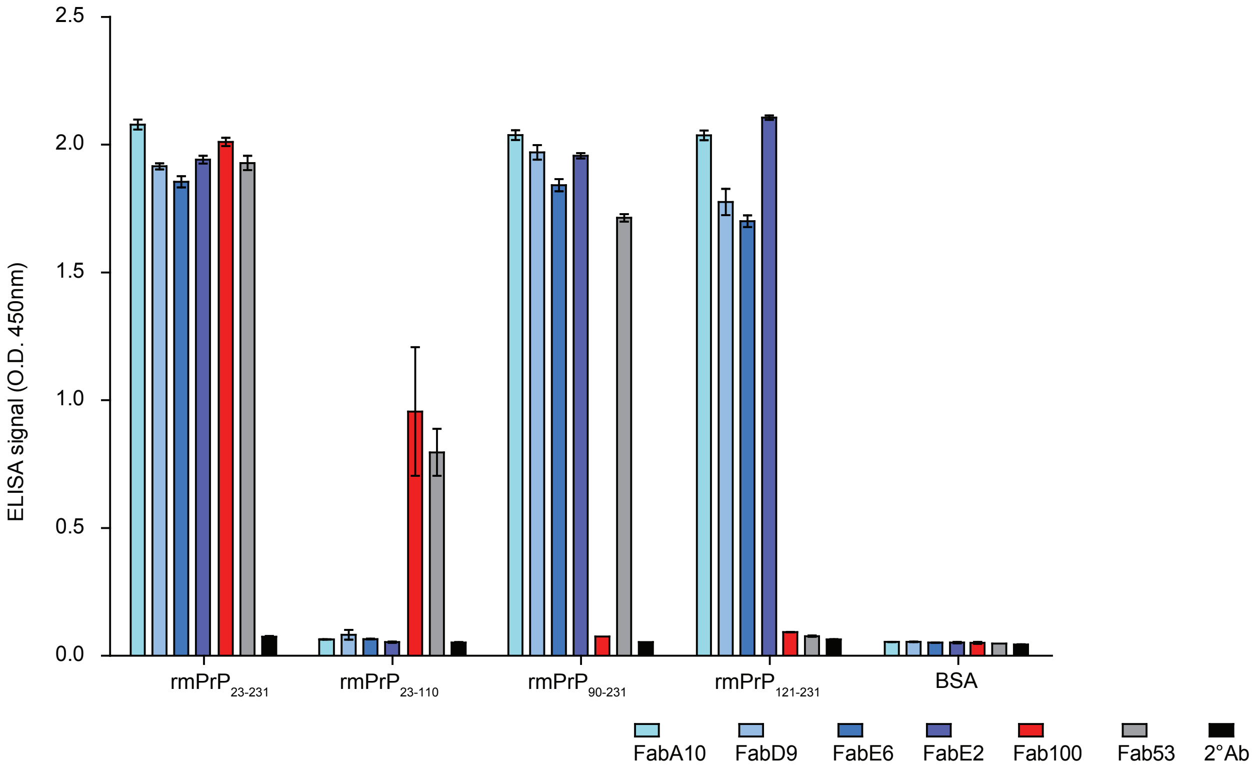


***Supplementary figure 18.*** ELISA (OD at 450nm) comparing the reactivity of phage-derived anti-PrP Fabs to full-length $rmPrP_{23\text{-}231}$, FT fragment $rmPrP_{23\text{-}110}$ and GD fragments $rmPrP_{90\text{-}231}$ and $rmPrP_{121\text{-}231}$. Anti-PrP Fab100 and Fab53 bind within the FT of PrP - the octapeptide repeat region (OR, amino acid 51-90) and the charged cluster 2 (CC2, amino acid 93-100), respectively. FabA10, FabD9, FabE6 and FabE2 bind within the GD.
Error bars = standard error of the mean.
