## Supplementary Figure S19 for "A conformational switch controlling the toxicity of the prion protein"

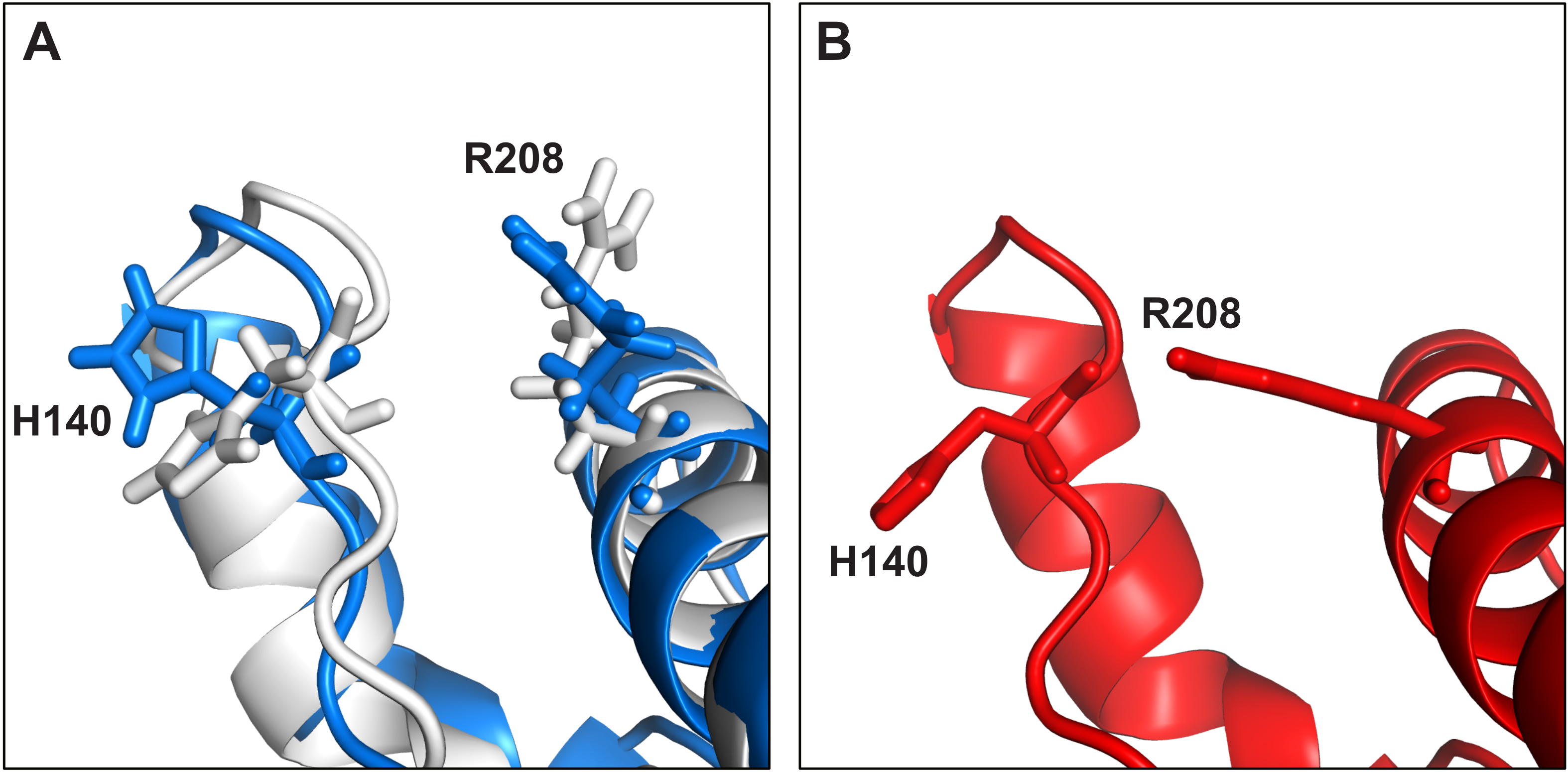


***Supplementary Figure 19.*** The R208-H140 interaction is present in POM1-bound PrP (**B**, *red*) but not in free PrP (**A**, *white*) or in its complex with FabA10 (**A**, *blue*). The final state of MD simulations starting from a POM1-bound conformation, with R208-H140 interaction present, is shown for FabA10.
