## Supplementary Figure S20 for "A conformational switch controlling the toxicity of the prion protein"

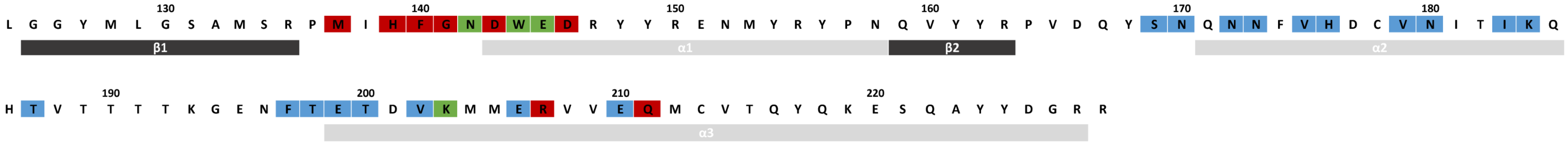


***Supplementary Figure 20.*** Overlap (green) of FabA10 (blue) and POM1 (red) epitopes on murine PrP$^{C}$-GD. Coloring according to Fig. 4D+E.
